# New insights into the evolution of SPX gene family from algae to legumes; a focus on soybean

**DOI:** 10.1101/2021.08.24.457498

**Authors:** Mahnaz Nezamivand Chegini, Esmaeil Ebrahimie, Ahmad Tahmasebi, Ali Moghadam, Saied Eshghi, Manijeh Mohammadi-Dehchesmeh, Stanislav Kopriva, Ali Niazi

## Abstract

**Background:** SPX-containing proteins have been known as key players in phosphate signaling and homeostasis. In Arabidopsis and rice, functions of some SPXs have been characterized, but little is known about their function in other plants, especially in the legumes.

**Results:** We analyzed SPX gene family evolution in legumes and in a number of key species from algae to angiosperms. We found that SPX harboring proteins showed fluctuations in domain fusions from algae to the angiosperms with, finally, four classes appearing and being retained in the land plants. Despite these fluctuations, Lysine Surface Cluster (KSC), and the third residue of Phosphate Binding Sites (PBS) showed complete conservation in almost all of SPXs except few proteins in *Selaginella moellendorffii* and *Papaver sumniferum,* suggesting they might have different ligand preferences. In addition, we found that the WGD/segmentally or dispersed duplication types were the most frequent contributors to the SPX expansion, and that there is a positive correlation between the amount of WGD contribution to the SPX expansion in individual species and its number of EXS genes. We could also reveal that except SPX class genes, other classes lost the collinearity relationships among Arabidopsis and legume genomes. The sub- or neo-functionalization of the duplicated genes in the legumes makes it difficult to find the functional orthologous genes. Therefore, we used two different methods to identify functional orthologs in soybean and Medicago. High variance in the dynamic and spatial expression pattern of GmSPXs proved the new or sub-functionalization in the paralogs.

**Conclusion:** This comprehensive analysis revealed how SPX gene family evolved from algae to legumes and also discovered several new domains fused to SPX domain in algae. In addition, we hypothesized that there different phosphate sensing mechanisms might occur in *S. moellendorffii* and *P. sumniferum*. Finally, we predicted putative functional orthologs of AtSPXs in the legumes, especially, orthologs of AtPHO1 and AtPHO1;H1, involved in long-distance Pi transportation. These findings help to understand evolution of phosphate signaling and might underpin development of new legume varieties with improved phosphate use efficiency.

## Background

Phosphorus (P) as an essential macronutrient serves as a structural element for many organic compounds, involved in multiple biosynthetic and metabolic processes [1, 2]. P containing molecules play a central role in various physiological processes, including respiration, photosynthesis, membrane transport, regulation of enzyme activity, oxidation-reduction reactions and signal transduction throughout plant growth, and development [3, 4]. Therefore, plants have evolved a number of mechanisms to ensure that P is readily available for all these processes. In particular, a wide range of responses are induced by phosphate (Pi) starvation [5, 6]. The regulation occurs at both transcriptional and posttranscriptional levels and many components of the regulatory network are known. The central regulator of the Pi starvation response and signaling network is the MYB transcription factor, AtPHR1 or OsPHR2 [7–9]. The PHR factors are negatively regulated through interaction with SPX domain proteins, which serve as sensors of P-status of the cells. In high P availability, inositol polyphosphates (PP-InsPs) bind to the basic surface of SPX domain proteins and facilitate their binding to PHR. This interaction may sequester PHR1 in the cytosol or prevent its association with DNA in the nucleus [10]. In low P supply, low availability of PP-InsPs-SPX results in the release of PHR1 to translocate to nucleus and to activate Pi starvation induced (PSI) genes [8]. Additionally, SPX domain proteins were shown to be involved in nitrate-phosphate signaling crosstalk in rice where nitrate-dependent interaction with NRT1.1B caused ubiquitination and degradation of OsSPX4 and consequently translocation of OsPHR2 and OsNLP3 into nucleus to induce PSI genes and nitrate inducible genes, respectively [11].

Despite the importance of SPX domain proteins in Pi signaling and nitrogen-dependent phosphate homeostasis, the functionality of all these proteins is still unclear. SPX domain proteins are important components of plant Pi homeostasis and can be divided into four classes based on the presence of extra domains: while class 1 only includes SPX domain, other three classes (SPX-EXS, SPX-MFS, SPX-RING), contain extra EXS, MFS, or RING domains, respectively [6]. There are four and six members of the SPX class 1 in Arabidopsis and rice, respectively [12, 13] as AtSPX3 and OsSPX1 act as negative regulators of Pi starvation signaling [12, 13]. Indeed, AtSPX1, localized in the nucleus, has a high binding affinity for AtPHR1 under high P condition and prevents it from activation of the downstream Pi starvation-induced (PSI) genes [8]. The rice OsSPX4 protein involved in the nitrate dependent regulation of Pi uptake [11] also belongs to this class. The most functional variation was observed in the EXS class members, including AtPHO1 and AtPHO1;1 involved in long-distance Pi transport from roots to shoots [14, 15], AtPHO1;4 with a role in response of hypocotyls to blue light [16], seed size and flowering [17–19] and AtPHO1;10 being induced by numerous stresses, such as local wounding [20, 21]. The Major Facilitator Superfamily (MFS) domain confers transport activity, therefore, SPX-MFS class are involved in both transport and signaling [22]. Finally, members of SPX-RING class are also called Nitrogen Limitation Adaptation (NLA) proteins due to their first identified role in nitrogen starvation resistance [23].

Recently, two other classes of SPX proteins, SPX-SLC and SPX-VTC, were characterized in algae as involved in polyphosphate synthesis and its transportation into vacuoles [24]. These two classes seem to be lost during the evolution of plants with shifting the type of phosphate storage from polyP in algae to Pi in the later-diverging Streptophytes [24]. It seems there have been some extra domains fused with SPX domain that might have been lost during the evolution of SPX proteins and that have not been comprehensively explored yet [25].

Legumes (Fabaceae) are the second most important family of crop plants economically [26]. Characterization of the SPX gene family in legumes can be helpful to gain insights into mechanisms of Pi homeostasis and thus underpin development of P efficient varieties. In this study, we performed a comprehensive analysis of SPX proteins from several legume crops (soybean, alfalfa, and common bean), and compared with species of more basal taxonomic groups such as mosses (*Phiscomitrella patens*), liverworts (*Marchantia polymorpha*), lycophytes (*Selaginella moellendorffii*), basal angiosperms (*Papaver somniferum*, *Amborella trichopoda*, and *Nymphaea colorata*), Rhodophytes (*Cyanidioschyzon merolae*, *Galdieria sulphuraria*, and *Chondrus crispus*), chlorophytes (*Chlamydomonas reinhardtii* and *Ostreococcus lucimarinus*), and charophytes (*Chara braunii*). We analyzed SPX protein evolution through phylogenetic analysis, conserved motif changes, and identification of ancestral motifs. In addition, because of only a partial functional characterization of SPX in legumes [27–31], we identified their functional orthologs with well-characterized SPXs from *Arabidopsis thaliana*. Since sequence-based orthology identifications alone have weakness in the one-to-many or many-to-many orthologs, expressologs identification was used as a complementary approach for functional ortholog identification [32]. With the combination of these two methods, we identified the functional orthologs of key regulators AtPHO1, AtPHO1;H1, AtSPX4, AtPHO1;H10, and AtNLA2 in the three legumes. In addition, we identified novel domains in SPX proteins of algae and functionally characterized SPX proteins in soybean and Medicago.

## Results

### Identification of SPX domain proteins from algae to legumes

While in several plant species four families of SPX proteins were characterized, much less is known about these proteins in legumes: in soybean and common bean just 10 and 3 members of class 1 were characterized and no SPX proteins in *M. truncatula*. Therefore, we intended to characterize this protein family in these legume species and set it into evolutionary context by analysis of SPX proteins from algae and basal plants. Sequences of SPX proteins were obtained by BLASTP searches at EnsemblPlants from the legumes (*G. max*, *P. vulgaris*, and *M. truncatula*), moss (*P. patens*), liverwort (*M. polymorpha*), lycophyte (*S. moellendorffii*), basal angiosperms (*P. somniferum*, *A. trichopoda*, and *N. colorata*) rhodophytes (*C. merolae*, *G. sulphuraria*, and *C. crispus*), chlorophytes (*C. reinhardtii* and *O. lucimarinus*), and charophytes (*C. braunii*) protein databases using full-length amino acid sequences of SPXs from Arabidopsis (20 proteins). After removing sequences lacking the SPX domains and redundant and partial sequences, we compiled all SPX proteins in the latest version of protein database in EnsemblPlants for these 15 species. Some proteins were shorter than 200 aa and were excluded from further analyses, including four short proteins of soybean (*GLYMA_12G154800*, *GLYMA_10G097000*, *GLYMA_09G098200*, *GLYMA_20G032200)*, two partial proteins of common bean (*PHAVU_010G0720001g*, *PHAVU_010G0720000g*), one protein of *M. truncatula* (MTR_8g058603). In addition, we excluded one protein of *M. truncatula* (*MTR_0262S0060)*, where its corresponding gene is located on a scaffold but not chromosomes, and one protein of common bean (*PHAVU_007g1245000g*), which had different structure from other SPX genes. Finally, 34 SPX proteins in *G. max*, 19 in *M. truncatula*, 17 in *P. vulgaris*, 22 in *P. patens*, 10 in *M. polymorpha*, 2 in *C. merolae*, 4 in *G. sulphuraria*, 2 in *C. crispus*, 5 in *C. reinhardtii*, 2 in *O. lucimarinus*, 4 in *C. braunii*, 42 in *P. somniferum*, 11 in *A. trichopoda*, 16 in *N. colorata*, and 31 in *S. moellendorffii* were identified (Supplemental Table S1). Furthermore, the proteins were classified into the four subfamilies based on their additional domains. Interestingly, in some algae and basal plants, we found extra domains that have not been previously reported (Figure 1). Totally, among these species, class EXS was with 88 proteins the largest, followed by SPX class with 48 proteins and MFS and RING classes containing 29 and 26 proteins, respectively. Subsequently, the corresponding SPX genes in soybean, *M. truncatula* and common bean were named in each subfamily based on their chromosomal positions (Supplemental Table S1).

**Figure 1.**
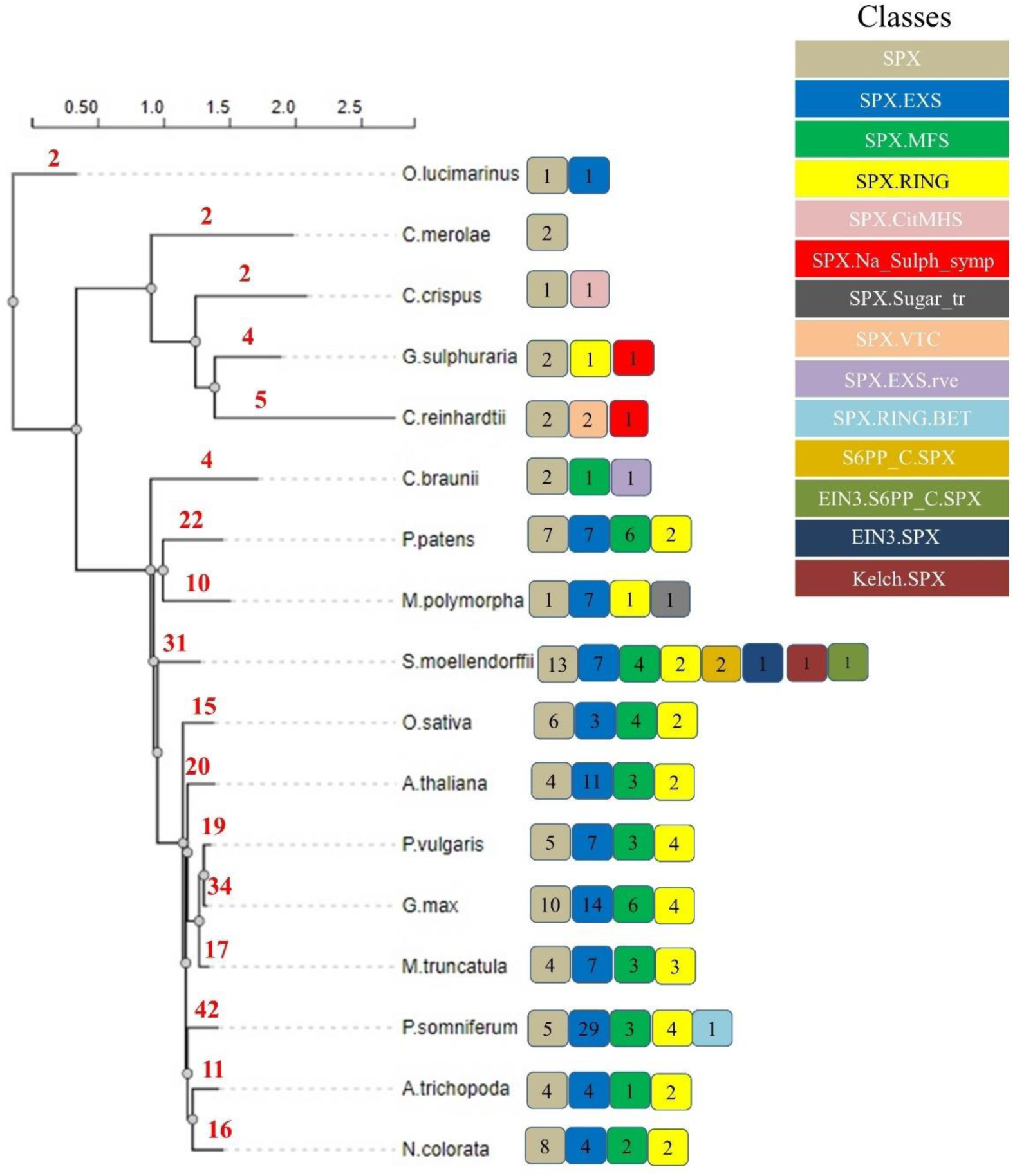
Evolution and frequency of genes in different SPX classes from algae to current Angiosperms. The species tree was constructed based on protein sequences of identified SPXs. Types of classes are shown in different colored boxes, the numbers in boxes represent the number of identified genes in each class while the total number of identified SPXs in each species is written in red on the branches.

As can be seen in the Figure 1, all basal and current angiosperms possess only the four main classes of SPX proteins. On the other hand, some additional domains were observed in liverwort, lycophyte, and algae based on Pfam and CDD scanning of sequences; SPX-VTC (vacuolar transporter chaperone), EIN3-SPX (Ethylene intensive 3), SPX-CitMHS (Citrate transporter), SPX-Na_sulph_symp (sodium sulphate symporter), SPX-RING-BET (Bromodomain extra-terminal-transcription regulation), S6PP_C-SPX (Sucrose-6F-phosphate phosphohydrolase C-terminal), EIN3-S6PP_C-SPX, Kelch-SPX (Galactose oxidase), SPX-EXS-rve, and SPX-Sugar_tr (Figure 1). The exact roles of these additional domains in the basal plants and algae are not completely known. It was previously reported that in some SPX proteins, SPX domain was located at C terminal instead of N terminal [33]. Indeed, we observed this structure in 4 different classes in *S. moellendorffii*, including EIN3-S6PP_C-SPX, Kelch-SPX, EIN3-SPX, and S6PP_C-SPX.

Predicted physiochemical and biochemical parameters of these SPX proteins in legume crops are listed in Supplemental Table S1. Indeed, members of the same subfamily have similar properties. The most variation in physiochemical parameters was observed in EXS class, while MFS class was the most similar. For example, lengths of all SPX-MFS proteins in the three species ranged from 691 to 700 aa, but the corresponding SPX-EXS proteins ranged from 475 to 1570 aa with the MtEXSs having the largest proteins in comparison with soybean and common bean. SPX-EXS and SPX-RING classes have the highest isoelectric point (pI), above 9 and 8, respectively. The calculated values for aliphatic index of SPX proteins show that the SPX-MFS subfamily have most thermostability, with a range of 105 to 111. GRAVY value (grand average of hydropathicity) is the sum of the hydropathy values of all amino acids divided by the protein length. Except for the proteins in the SPX-MFS subfamily, nearly all of the GmSPXs are hydrophilic, with a GRAVY value less than 0. Subcellular localization prediction performed with Wolf PSORT revealed that most of the GmSPX proteins are located in the plasma membrane or endomembrane system, followed by nucleus and chloroplast. In PSORT results, all members of SPX-EXS and SPX-MFS subfamilies were located in the plasma membrane, and all members of SPX-RING were located in nucleus, corresponding to the known functions of representatives of these subfamilies in Arabidopsis.

### Phylogenetic tree

Multiple alignment of the SPX protein sequences from soybean, *M. truncatula*, common bean, Arabidopsis, rice, wheat, rapeseed, *A. trichopoda*, *C. braunii*, *C. reinhardtii*, *C. crispus*, *C. merolae*, *G. sulphuraria*, *M. polymorpha*, *N. colorata*, *O. lucimarinus*, *P. somniferum*, *P. patens*, and *S. moellendorffii*, as well as proteins from mouse, human, and *Caenorhabditis elegans* as an out-group, followed by phylogenetic analysis revealed four distinct clades of SPX proteins, SPX, EXS, MFS, and RING (Figure 2). This topology and distinct separation of four classes are consistent with previous studies on SPX gene family [3, 12, 13, 27, 34]. SPX and EXS sequences formed two distinct clades, while MFS and RING along with box. C (*OSTLU26654.EXS*, *CHC T00007225001.SPX.CitMHS*, *CHLRE 09g251650V5.SPX.Na_Sulph_symp*, *C5167 020395.NLA.BET*, *Gsu16460.SPX.NLA*, and *CMP022C.SPX*) and box. D (*Gsu35240*.*SPX.Na_sulph_symp*) have diverged from a common ancestor and form the third major clade. SPX clade was divided into three sub-clades; SPX-I, SPX-II, and SPX-III. SPX-II and SPX-III are specific to the basal and current angiosperms and the proteins in these two sub-clades are homologs of AtSPX3 and ATSPX1/2, respectively. On the other hand, SPX-I is comprised from homologs of the basal plants (lycophytes, liverwort, moss) and algae and few proteins from the basal and current angiosperms, all being homologs of AtSPX4. Proteins in box A and in box B could be ancient homologs for SPX-I and SPX-II/III, respectively. Likewise, EXS clade was divided into three sub-clades; EXS-I is specific to lower plants (*S. moellendorffii, M. polymorpha*, and *P. patens*), EXS-II is a mixed group from monocots, eudicots, and basal angiosperms, all homologs of AtPHO1 and AtPHO1;H1, and EXS-III contain eudicots and the basal angiosperms without any genes of monocots. The outgroup genes used in this study were grouped in box E clustered with EXS clade. Overall, topology of EXS class is consistent with He et al., (2013), in that basal plants (lycophytes and moss) EXS homologs were grouped separately from the angiosperms, and also with the previous reports on EXS genes that monocots only possess homologs for AtPHO1 and AtPHO1;H1 [6, 24, 35].

**Figure 2.**
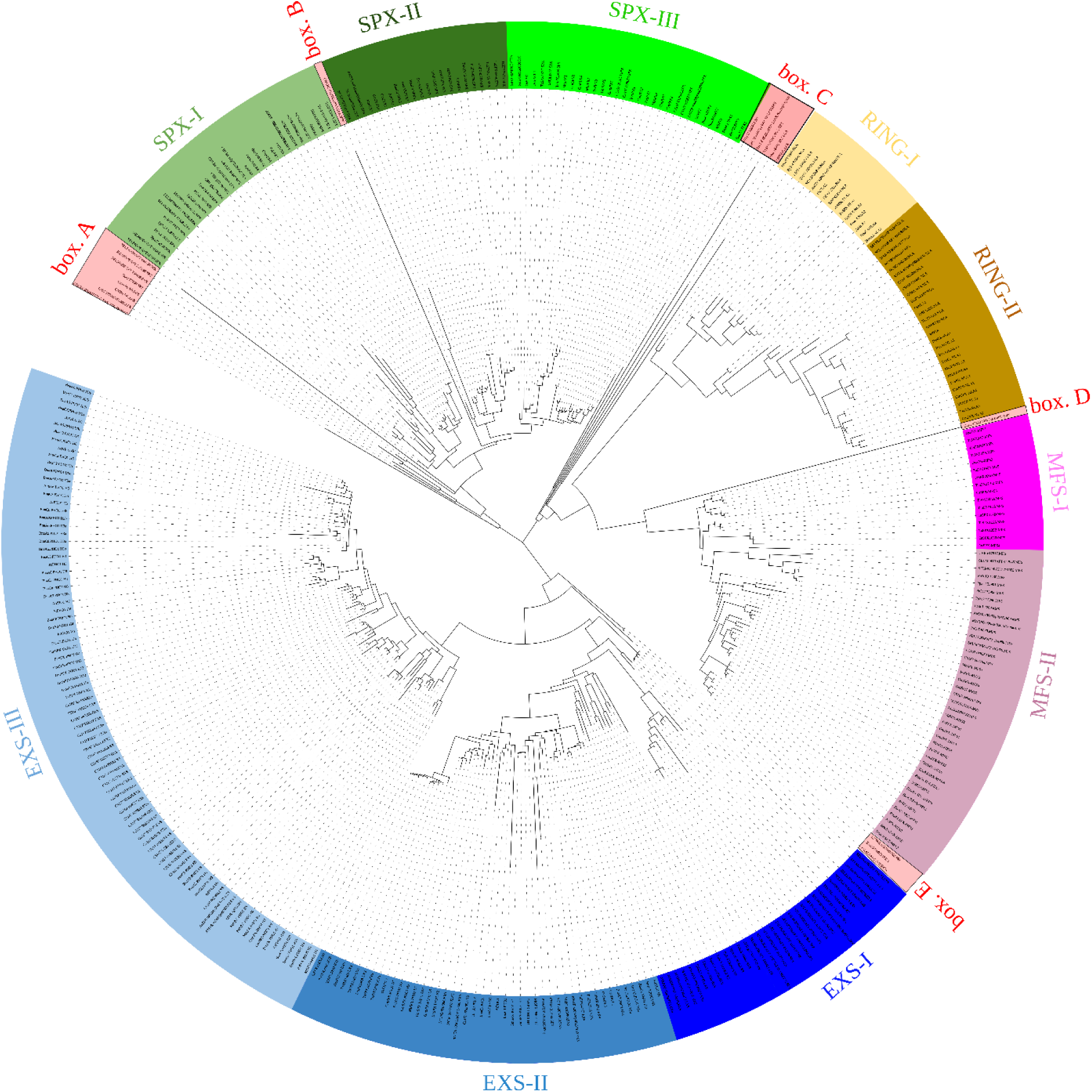
Phylogenetic analysis of 218 SPX containing proteins from 19 plant species. The phylogenetic tree was constructed using the Maximum Likelihood method. The SPX genes of Arabidopsis, rice, wheat, rapeseed, M. truncatula, soybean, and common bean are represented with At, Os, Ta, Bna, Mt, Gm, and Pv abbreviations, respectively. Other species are named based on their Gene IDs and their domains. Four different clades are marked in colors: SPX (green), RING (brown), MFS (pink), and EXS (blue). Sub-clades of each clade are shown with light and dark shades of the respective colors. Five boxes show paraphyletic branches; box E comprises the outgroup species.

Box C with ancient genes for both MFS and RING and box D as sister for MFS class together with MFS and RING clades seem to have evolved from a common ancestor. MFS homologs in monocots specifically grouped in MFS-I, while MFS-II contained all MFS orthologs from the other species. This could suggest that differentiation among MFS proteins has occurred after the divergence of monocot and dicots from a common ancestor. Similarly, RING clade was divided into two sub-clades, but both contained RING orthologs from all of species; RING-I was grouped with the ancestor from *P. patens*, while RING-II included *S. moellendorffii* orthologs as its sister. The overall tree topology is very similar to results of Wang et all (2021), who investigated SPX gene family in chlorophytes and streptophytes, with focus on algae.

### Protein motifs gain and loss in SPX family throughout evolution

Conserved protein motifs were predicted using MEME program for each SPX protein class and all species (Additional file 1: Figure S1 to S5). This analysis may explain when different classes of SPX proteins have appeared and how motifs were gained or lost in each class during the evolution. The ancestral motifs in SPX domains such as motifs 3, 4, 2, and 1 seem to originate from red algae (Additional file 1: Figure S1). There is a high fluctuation of motif composition during the evolution. Some motifs are species specific like motifs 13, 14, and 19 that are present only in legumes, probably arising after legume whole-genome duplication event. The most variability in the motif composition was observed in *S. moellendorffii* with some specific motifs like 8, 15, and 18. The lengths of proteins in angiosperms were very similar but shorter than in the basal plants. The EXS domain was detected only in *O. lucimarinus* with 9 motifs - 9, 6, 5, 2, 3, 11, 4, 10, and 1 (Additional file 1: Figure S2). Almost all these motifs have been retained during the evolution as ancestral motifs. In addition, some other motifs appeared in *C. braunii* such as 15, 7, 16, 20, 12, and 8, suggesting they were present in the common ancestor of Chlorophyta and Streptophyta. Although Wang et al [24] reported one SPX-MFS in *M. polymorpha* genome, we could not find an intact SPX-MFS domain, but SPX-Sugar_tr domain with a highly similar motif composition with other MFSs was identified (Additional file 1: Figure S3). As it has previously been reported, *PHT5* genes in *B. napus* have SPX domain connected to overlapping MFS and Sugar_tr domains [36], however, we only found SPX and Sugar-tr domains in *M. polymorpha* genome. The first SPX-MFS protein was observed in *C. braunii* with 18 common motifs with other species. Two newly observed motifs in *P. patens*, motifs 16 and 13, probably have evolved by dispersed duplication in *P. patens* and have been retained in all basal and current angiosperms. Interestingly, other five MFSs in *P. patens*, without the motifs 16 and 13, have been no longer found in angiosperms.

The evolutionary oldest NLA has been detected in *G. sulphuraria* and it was retained during the course of evolution of current angiosperms, but was not found in other Rhodophytes or Chlorophytes. In fact, the only NLA identified in *G. sulphuraria* just showed two motifs in common with other species, motifs 2 and 3 (Additional file 1: Figure S4). Therefore, these motifs could be considered as ancestral motifs of NLA class which then further evolved by dispersed duplication in *M. polymorpha*, adding motifs 8, 7, 1, and 6 into the ancestral domains. One NLA in *P. somniferum* underwent dispersed duplication and gained motif 10 that has only been retained in the core eudicots, while two NLAs in *S. moellendorffii* segmentally duplicated and gained two specific motifs 13 and 19. Motif 16 was just observed in legume genomes that might evolved after legume whole-genome duplication (WGD) event. The most variability in motif composition of NLA class was observed in *P. somniferum*. Motif compositions in the new identified classes showed a high variation and it was impossible to find their ancestral motif (Additional file 1: Figure S5). However, it could be concluded that SPX-Na_Sulph_sym and SPX-CitMHS with high similarity in the motif composition, probably have similar origin and function. In summary, during the evolution different duplication events added new motifs to the ancestral motifs and other motifs specifically appeared in individual species to acquire new functions.

### Consensus sequences of SPX domains from algae to eudicots

We then predicted conserved motifs among all identified SPXs (Additional file 1: Figure S6). There are four conserved motifs in SPX class members, among them two motifs, 2 and 4, are common in the almost whole span of SPXs. Therefore, we can hypothesize that these two motifs have an important role for all SPXs. Afterwards, consensus sequences of these two motifs were constructed across all phyla (algae, charophytes, liverwort, bryophytes, lycophytes, basal angiosperms, and current angiosperms) and also across each class (SPX, EXS, MFS, RING, and new identified classes) (Additional file 1: Figure S7-S10). Motif 4 is 29 aa in length and was present in all SPX proteins except the following ten: C5167_005902.EXS, C5167_032842.EXS, C5167_043562.EXS, C5167_043565.EXS, C5167_003186.NLA, C5167_046257.NLA, SELMODRAFT_419593.SPX, SELMODRAFT_419593.SPX, OsSPX4 and PvPHO1. Five amino acid residues, number 5, 9, 15, 19, and 24, were almost 100% conserved, except the fifth residue in *C. braunii* (Additional file 1: Figure S7). Regarding conservation in different classes (Additional file 1: Figure S8), the leucine (residue 9) was completely conserved in the EXS, MFS, RING, and new identified classes, then the phenylalanine (residue 19) was completely conserved in EXS and MFS classes, but SPX class had some members with different residues in these five positions with a very high overall conservation in this class. In addition, each class had other conserved residues, suggesting special functions.

Motif 2 is 21 aa long and was absent in CHLRE_02g111650v5.SPX, AMTR_s00106p00066860.SPX, NC1G0101580.SPX, C5167_011965.SPX, Gasu_57230.SPX, C5167_043539.EXS, SELMODRAFT_450458.EXS, SELMODRAFT_431864.SPX, SELMODRAFT_419593.SPX, and only one protein from the current angiosperm, PvPHO1;5, which is a partial protein. This motif exhibited more conserved residues at positions 1, 7, 8, 14, 15, 16, 17, 18, 20, and 21. Residue 17 was completely conserved in the all proteins containing motif 2 and residues 14, 18, and 21 were conserved in the all proteins except a few in *S. moellendorffii* and *P. sumniferum* showing different residues instead of lysine (Additional file 1: Figure S9). The lysine residues 14, 17, and 21 form a Lysine Surface Cluster (LSC), and were found to interact with sulfate in the crystal structure of human phosphate transporter XPR1, and to be a part of a larger binding site for PP-InsP [9]. Consequently, in the different classes of the SPX proteins (Additional file 1: Figure S10), some of the 10 conserved positions were completely conserved such as K1, N8, KILKK (14 to 18) in RING and MFS, K18 in SPX, K21 in RING, and N8, I15, K18, as well as K21 were completely conserved across the new identified classes. Overall, these two motifs were conserved in all but a few proteins from *S. moellendorffii* and *P. sumniferum*, PvPHO1;5, and OsSPX4, implying that they might possibly interact with InsP/PP-InsP in a different manner, as previously reported for OsSPX4 [9]. In addition, different conserved residues in different classes could suggest that they may have different phosphate-containing ligand or different levels of Pi in cells

### Expansion pattern of SPX genes and collinearity analysis

To pinpoint the expansion modes in the land plants, we investigated duplication types in basal and current angiosperms, liverwort, hornwort, and *S. moellendorffii* (Figure 3 and Supplemental Table S2). Taken together, WGD, segmental, and dispersed duplications contributed most to the SPX gene family expansion. The expansion patterns in soybean, *P. somniferum*, *N. colorota*, and *S. moellendorffii* mostly arose from WGD/segmental duplication type. However, *S. moellendorffii* did not have any WGD events, therefore, its expansion and unique SPX classes must have arisen through local or segmental gene duplication [37]. WGD/segmental duplication type did not participate in the SPX expansion in *A. trichopoda* and *M. polymorpha* genomes and it only resulted in one duplicated block in *P. patens* genome. In these three species, SPX expansion were affected mostly by dispersed duplication type. The high number of WGD/segmental types of duplication in *S. moellendorffii*, soybean, and *P. somniferum* can shed light on the reason of high variation of gene family sizes in the closely related plants.

**Figure 3.**
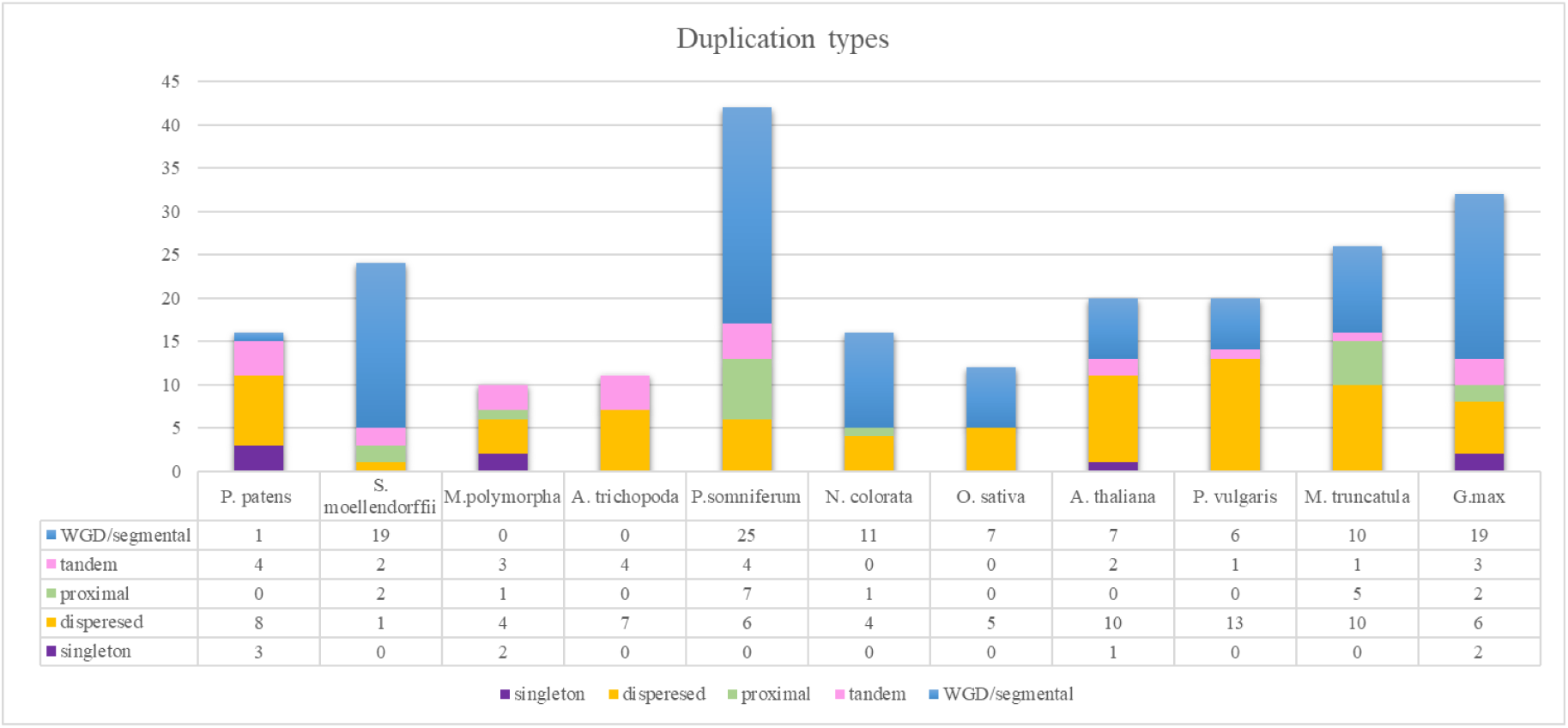
SPX gene family expansion from algae to the current Angiosperms. Duplication event types were predicted in the *P. patens*, *S. moellendorffii*, *M. polymorpha*, *A. trichopoda*, *P. somniferum*, *N. colorota*, *O. sativa*, *A. thaliana*, *P. vulgaris*, *M. truncatula*, *G. max*.

To get more information about evolutionary process of genes, collinearity analysis can provide information about conserved genomic regions of genes in different species [38]. Synteny relationship among two or a set of genes from two species means that they located in the same chromosome [39], but collinearity is a specific form of synteny with conserved gene order [40]. Collinearity analysis was conducted in three steps; 1. across *P. somniferum*, *N. colorota*, rice, Arabidopsis, and three legumes 2. Among *P. somniferum* and *N. colorota*, *P. patens*, and *S. moellendorffii* and 3. Among legumes. Collinearity analysis among legume crops, Arabidopsis, rice, and two basal angiosperms; *P. somniferum* and *N. colorata* discovered 121 collinear blocks (Figure 4, Supplemental Table S3); 30 blocks in Gm/Pv, 23 blocks in Gm/Mt, 15 blocks in Gm/Gm, 14 blocks in Ps/Ps, 10 blocks in Gm/At, 6 blocks in Nc/Nc and Pv/Mt, 3 blocks in Ps/Nc, Mt/At, Pv/At, and Os/Os, 2 blocks in Ps/Gm, and 1 block in At/At, Pv/Pv, Ps/Mt, and Mt/Mt. Rice as the only monocot in this analysis did not show any collinearity relationship for SPX gene family with other species.

**Figure 4.**
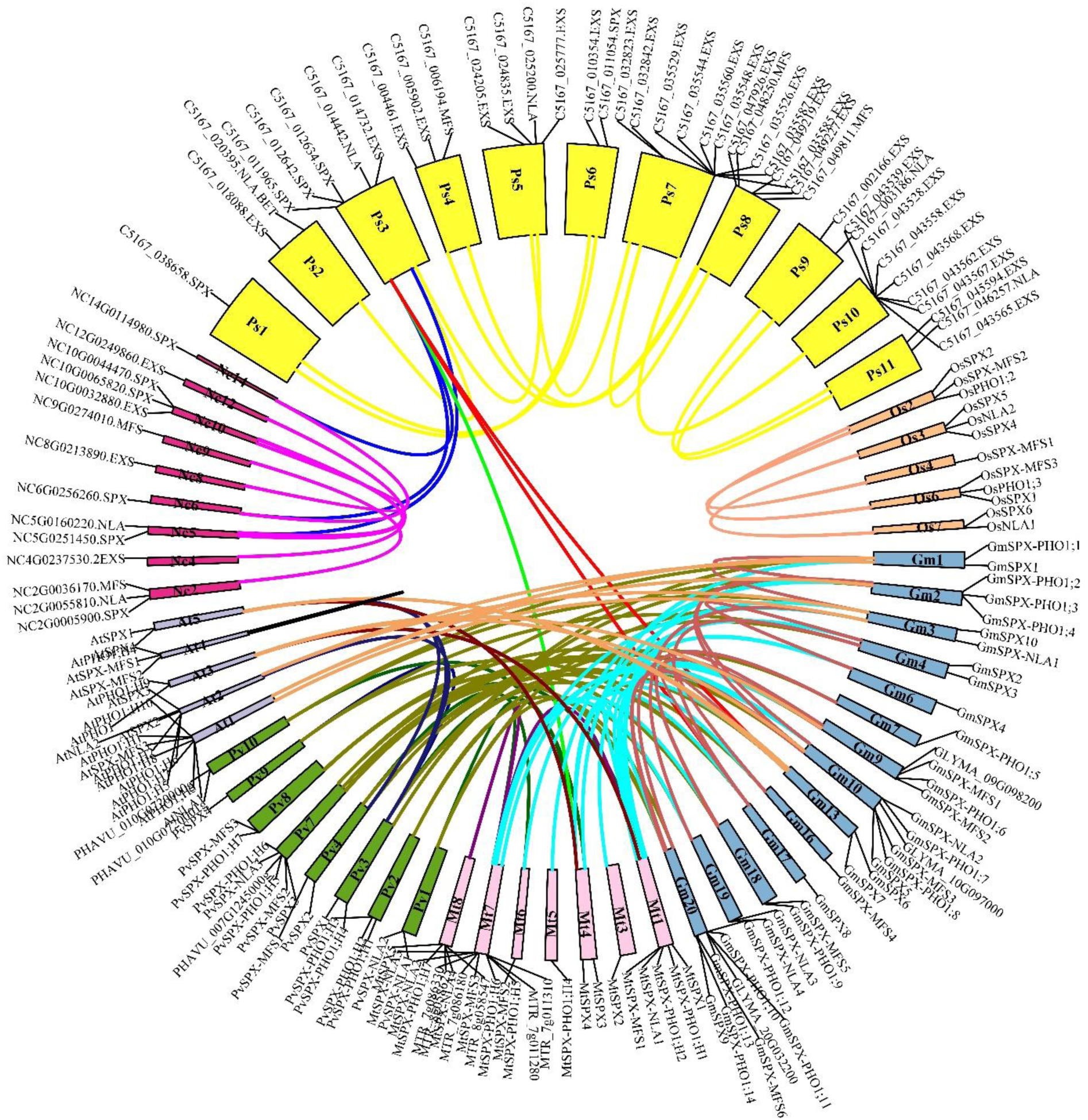
Circular collinearity plot of SPX gene family members among *G. max* (blue), *M. truncatula* (pink), *P. vulgaris* (green), *A. thaliana* (grey), *O. sativa* (orange), *P. somniferum* (yellow), and *N. colorota* (red). Collinear genes are linked by lines and boxes are representing chromosomes.

Collinear SPX genes among *P. somniferum*, *S. moellendorffii*, *N. colorata*, and *A. trichopoda* were predicted (Supplemental Table S3). *S. moellendorffii* did not show any collinearity relationship with other species, while *N. colorata* and *P. somniferum* had the most inter species collinear relationships (14). The most intra-genome collinear relationships were found in *P. somniferum* (14) and *S. moellendorffii* (10). The collinear analysis was performed also for the three legume crops (Figure 5, Supplemental Table S3). Of the 34, 19, and 17 SPX genes in soybean, *M. truncatula*, and common bean 32, 14, and 15 genes participated in collinear blocks. In total, 78 collinearity blocks between these plant species were discovered. A high level of collinearity relationships was found at 27/30 SPX genes in soybean/common bean and 19/23 SPX genes in soybean/*M. truncatula*, while the corresponding figure for *M. truncatula*/common bean was 6/7. However, just 15, 7, and 2 collinearity blocks were found in soybean/soybean, *M. truncatula* /*M. truncatula*, and common bean/common bean groups. All in all, after these three collinearity analyses, we concluded that inter-species collinearity patterns among basal angiosperms and among current angiosperms have changed. Across basal angiosperms, SPX class had the least inter-species collinearity, while among Arabidopsis and legumes, SPX showed the most inter-collinearity relationships. It can be concluded that except in SPX class, collinearity in the other classes has been lost.

**Figure 5.**
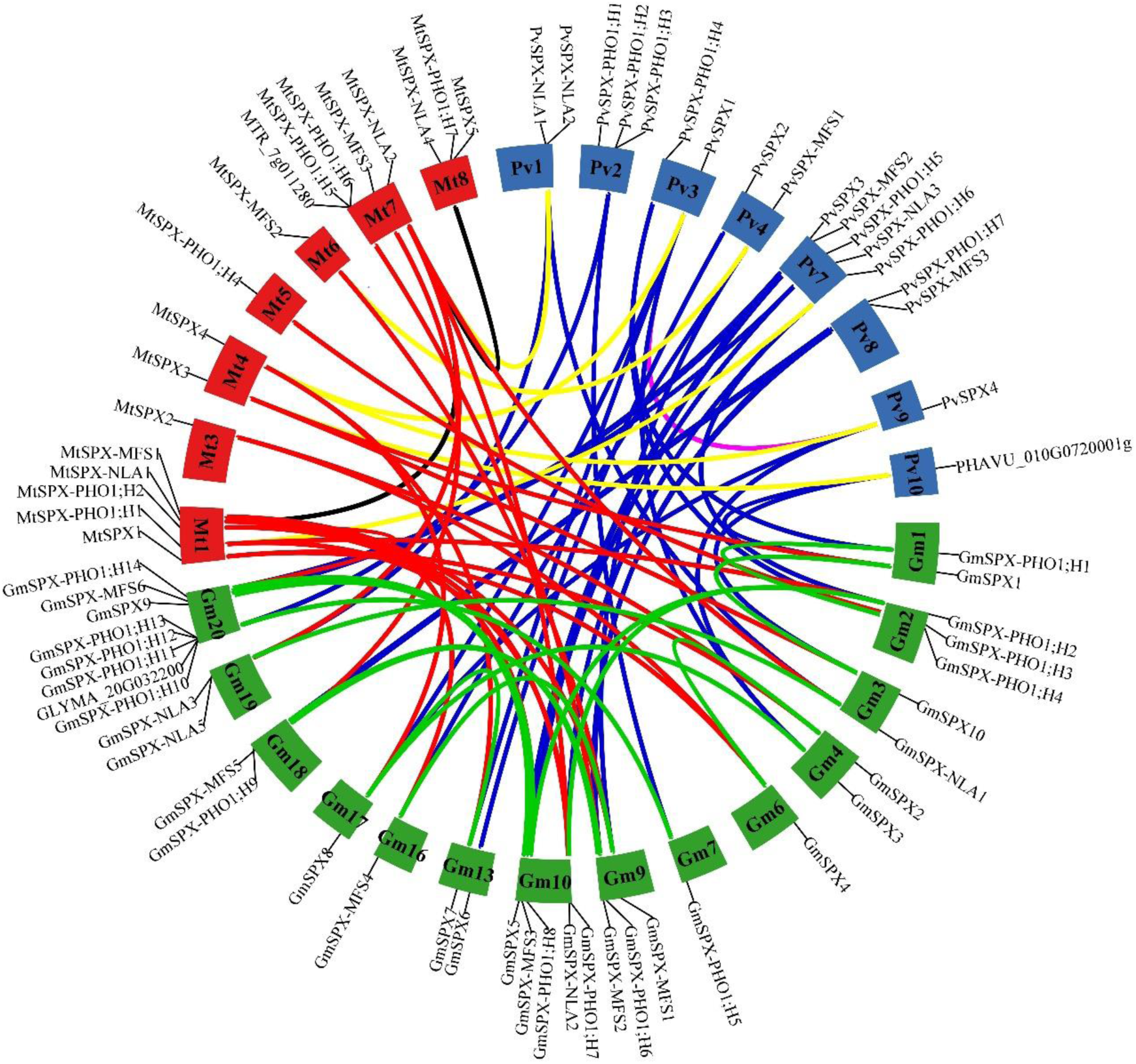
Circular collinearity plot of SPX gene family members among *G. max*, *M. truncatula*, *P. vulgaris*. Chromosomes of *G. max*, *M. truncatula* and *P. vulgaris* are respectively in green, red and blue. Links between *G. max* and *M. truncatula* are colored red, *G. max* and *P. vulgaris* in blue, *M. truncatula* and *P. vulgaris* in yellow as well as links within *G. max*, *M. truncatula* and *P. vulgaris* are colored in green, black and pink.

### Evolution of Cis-acting elements from algae to eudicots

Transcription factors bind to the cis-acting elements (CREs) in the promoter and regulate the transcription of corresponding genes [41]. Therefore, genes with similar expression patterns may contain the same regulatory elements in their promoters [27]. To explore whether transcription factor biding sites have evolved together with the coding regions of SPX genes, 1.5 kb upstream of the transcriptional start sites of all identified SPXs were downloaded and analyzed using PlantCARE database. In total, 124 CREs were detected (Supplemental Table S4) that can be classified in three major groups: responsive to abiotic stresses (drought, low temperature, hypoxia, wounding, defense, and stress), hormones (gibberellin, abscisic acid (ABA), salicylic acid (SA), ethylene, methyl jasmonate (MeJA), and auxin), and development-related elements (endosperm, meristem, MYB, and zein metabolism regulation). After the essential elements in promoter like TATA-box and CAAT-box, the most highly represented cis-acting elements were those involved in response to MeJA (CGTCA-motif and TGAG-motif) and ABA (ABRE and ARE). Looking for evolutionary pattern in these cis-acting elements, we performed hierarchical clustering on principal components (HCPC) using FactMineR-package. The HCPC grouped the genes into three clusters (Additional file 1: Figure S11).

Almost all SPXs from the current angiosperms fell into cluster 1 along with 2 SPXs of *G. sulphuraria* and few SPXs from basal angiosperms (Table 1, Additional file 1: Figure S11). Cluster 2 comprised mostly genes from basal angiosperms and few members of the current angiosperms, as well as all SPXs of *C. reinhardtii* and two SPXs from *G. sulphuraria*. Cluster 3, the smallest cluster, had 12 genes mostly from *P. patens* and just one SPX of the current angiosperms, *AtPHO1;H5*. Trying to find an evolutionary pattern across these clusters, we found out that they showed different frequencies of two MeJA responsive elements, TGACG and CGTGA motifs, that in cluster 3 all genes, in cluster 2 around 93%, and in cluster 1 only around 62% of genes possessed these two elements (Table 1). Besides, we extracted the most enriched CREs in each cluster to visualize frequencies of these elements across clusters. As can be seen in the Additional file 1: Figure S12, CREs involved in the developmental processes (CCGTCC motif, CCGTCC box, A-box) and stress response (DRE core, MYB recognition site, CCAT box) were significantly higher in cluster 2 than in the other clusters. Cluster 1 had higher frequency of two hormone responsive elements, TCA (salicylic acid responsive elements) and ERE (Ethylene-responsive elements) in comparison to the other clusters. Overall, it seems that during the evolution of angiosperms, SPX promoters were enriched by stress responsive elements and hormonal responsive elements, especially ERE and TCA.

**Table 1.** Number of genes having MeJA and ABA responsiveness elements in their promoter sequence.

| Clusters | TGACG-motif | CGTCA-motif | ABRE | ARE | Number of SPXs of<br>each species | Total |
| --- | --- | --- | --- | --- | --- | --- |
| Cluster 1 | 57 | 57 | 59 | 72 | 16 At, 27 Gm, 12 Mt,<br>15 Pv, 11 Pp, 1 CHb, 2<br>Gsu, 4 Nc, 2 Pp, 1 Mp, | 91 |
| Cluster 2 | 55 | 55 | 32 | 47 | 5 CHLRE, 2Gsu, 7Mp,<br>13 Ps, 2 CHb, 7 Nc, 12<br>Pp, 3 Sm, 2 Mt, 3 Gm, 3<br>At | 59 |
| Cluster 3 | 12 | 12 | 11 | 12 | 3 Pp, 1 Mp, 1 Sm, 1 At,<br>6 Ps, | 12 |

### Selective pressure and SPX history model in legumes

The Ks (number of synonymous substitutions per synonymous site) and Ka (number of nonsynonymous substitutions per nonsynonymous site) values of pairs of segmental duplicated SPX genes in soybean, *M. truncatula* and common bean were retrieved from Plant Genome Duplication Database (PGDD) (Supplemental Table S5). The Ka/Ks ratios < 1 indicate purifying selection and Ka/Ks values > 1 indicate positive selection [42, 43]. The Ka/Ks values for all pairs of segmental duplicated genes were < 0.3 implying an intense purifying selection on these gene pairs (Supplemental Table S5). In addition, the Ka/Ks ratio of duplicated gene pairs between soybean and *M. truncatula*, soybean and common bean, and *M. truncatula* and common bean were retrieved (Supplemental Table S5). The mean Ka/Ks values of 0.18, 0.16, and 0.14, respectively, suggest that the genetic pairs between species were subjected to purifying selection.

Based on the Ks values of duplication blocks retrieved from PGDD, the divergence times were estimated. In total, 36, 7, and 3 duplication blocks were retrieved for soybean, *M. truncatula*, and common bean, respectively (Supplemental Table S5). All duplication blocks related to MFS and RING class have Ks < 1.5, and the most recent duplication events belonged to MFS members in soybean. Evolutionary process of GmSPX genes was modeled based on Ks of duplication blocks (Figure 6). The duplicated SPX genes in SPX, EXS, MFS, and RING were classified into 3, 2, 2, and 1 groups, respectively. GmSPX-A firstly generated three copies after the Gamma WGT event, followed by loss of one copy. The two retained copies were further doubled after Legume WGD event, and after losing one copy, the rest three copies duplicated after Glycine WGD event, resulting in genes, GmSPX8, GmSPX7, GmSPX3, GmSPX4, and GmSPX2. GmSPX3 lost its linked duplicated gene (Figure 6). Unexpectedly, all three generated copies of GmSPX-EXS-A in Gamma WGT event were retained but their duplicated genes after Legume WGD were lost. Therefore, Glycine WGD resulted in generation of five genes (*GmSPX-PHO1;10*, *GmSPX-PHO1;5*, *GmSPX-PHO1;3*, *GmSPX-PHO1;9*, and *GmSPX-PHO1;6)* after a loss of one of the linked genes. However, GmSPX-EXS-B lost one copy in the first and second round of duplication events and lastly generated six genes (*GmSPX-PHO1;1, GmSPX-PHO1;4, GmSPX-PHO1;14, GmSPX-PHO1;8, GmSPX-PHO1;7, GmSPX-PHO1;2)*. *GmSPX-B* and -*C* as well as *GmSPX-MFS-B* shared the same evolutionary trajectory and generated two duplicated genes in the same way after three rounds of the evolution processes. In addition, *GmSPX-MFS-A* and *GmSPX-RING* were somewhat similar as both produced two duplicated blocks, although one copy was lost in *GmSPX-RING*, resulting finally in three and four genes, respectively.

**Figure 6.**
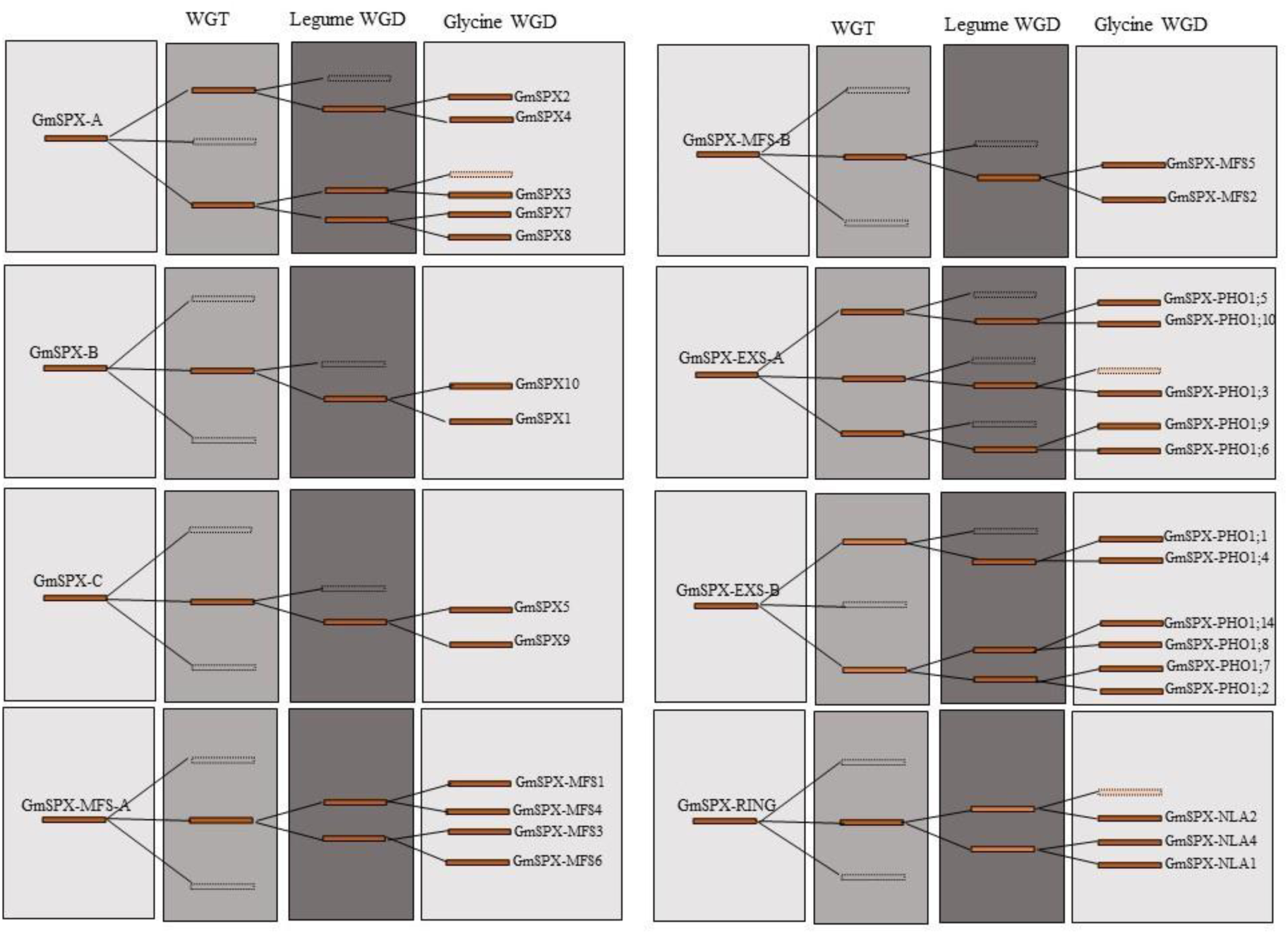
The evolutionary history of GmSPX genes. The reserved and lost blocks in the corresponding evolution are displayed by solid and empty blocks, respectively.

### Functional characterization of orthologous genes in legumes

Orthologs and orthogroups among seven current angiosperms were determined with OrthoFinder. Altogether, from 218 genes, 216 genes could be classified in seven orthogroups and just two genes of rapeseed (*BnaA6.PHO1;H3c* and *BnaA9.PHO1;H3b*) were not grouped, maybe suggesting a brassica-specific function for these proteins. All members of SPX, SPX-MFS, and SPX-RING were assigned into one group; 1, 3, and 4, respectively. On the other hand, members of EXS family were divided into four distinct groups: group 2 that was dicot-specific; group 7, brassicaceae-specific; as well as groups 5 and 6 that contained genes from all species (Table 2). All genes in an orthogroup are descended from a single ancestral gene.

**Table 2.** Ortholog groups among soybean, common bean, Medicago, Arabidopsis, rice, wheat, and brassica.

| Orthogroup | Pv | Gm | Ath | Bna | Os | Ta | Mt | Total |  |
| --- | --- | --- | --- | --- | --- | --- | --- | --- | --- |
| 1 | 4 | 10 | 4 | 11 | 5 | 15 | 5 | 54 | SPX group |
| 2 | 4 | 8 | 8 | 29 | 0 | 0 | 4 | 53 | SPX.EXS dicot-specific group |
| 3 | 3 | 6 | 3 | 8 | 4 | 12 | 3 | 39 | SPX.MFS group |
| 4 | 3 | 4 | 2 | 7 | 2 | 7 | 4 | 29 | SPX.RING group |
| 5 | 2 | 4 | 1 | 4 | 1 | 9 | 2 | 23 | SPX.EXS group |
| 6 | 1 | 2 | 1 | 5 | 2 | 3 | 1 | 15 | SPX.EXS group |
| 7 | 0 | 0 | 1 | 2 | 0 | 0 | 0 | 3 | SPX.EXS brassicaceae-specific group |

Orthologous genes across Arabidopsis and the three legume crops are presented in Table 3. Some genes showed a simple one-to-one orthology relationship, such as *GmSPX6*, *PvSPX2*, and *MtSPX5* with *AtSPX4*; *GmPHO1;3*, *PvPHO1;1*, and *MtPHO1;4* with *AtPHO1;H10*; and *GmNLA3*, *PvNLA1*, and *MtNLA2* with *AtNLA2*. Others showed one-to-many and many-to-many orthology relationships. Interestingly, the pattern of *AtSPXs* orthology relationships were the same among three legumes, and each SPX gene has the same evolutionary trajectories. To overcome the difficulty of one-to-many and many-to-many orthology inference, expressologs of *AtSPXs* with soybean and Medicago were retrieved from the Expression Tree Viewer [32]. Expression Tree Viewer allows to visualize expressologs depending on both sequence similarity and expression pattern similarity. Implementing this web tool resulted in postulating expressologs between Arabidopsis and soybean and Medicago (Supplemental Table S6). Generally, the results were in very good agreement with previous results from phylogenetic tree and OrthoFinder. Based on the Expression Tree Viewer results, we could designate *GmPHO1;2/7* and *MtPHO1;1/2* as the functional orthologs of *AtPHO1* and *AtPHO1;H1* with the function of long-distance Pi transport. However, it was difficult to find expressologs for other SPXs. Consistently, the function of *GmSPX1* [31] and *GmSPX3* [29] were characterized with negative and positive regulatory roles in phosphate deficiency that are the same for *AtSPX1/2* and *AtSPX3* [6].

**Table 3.**
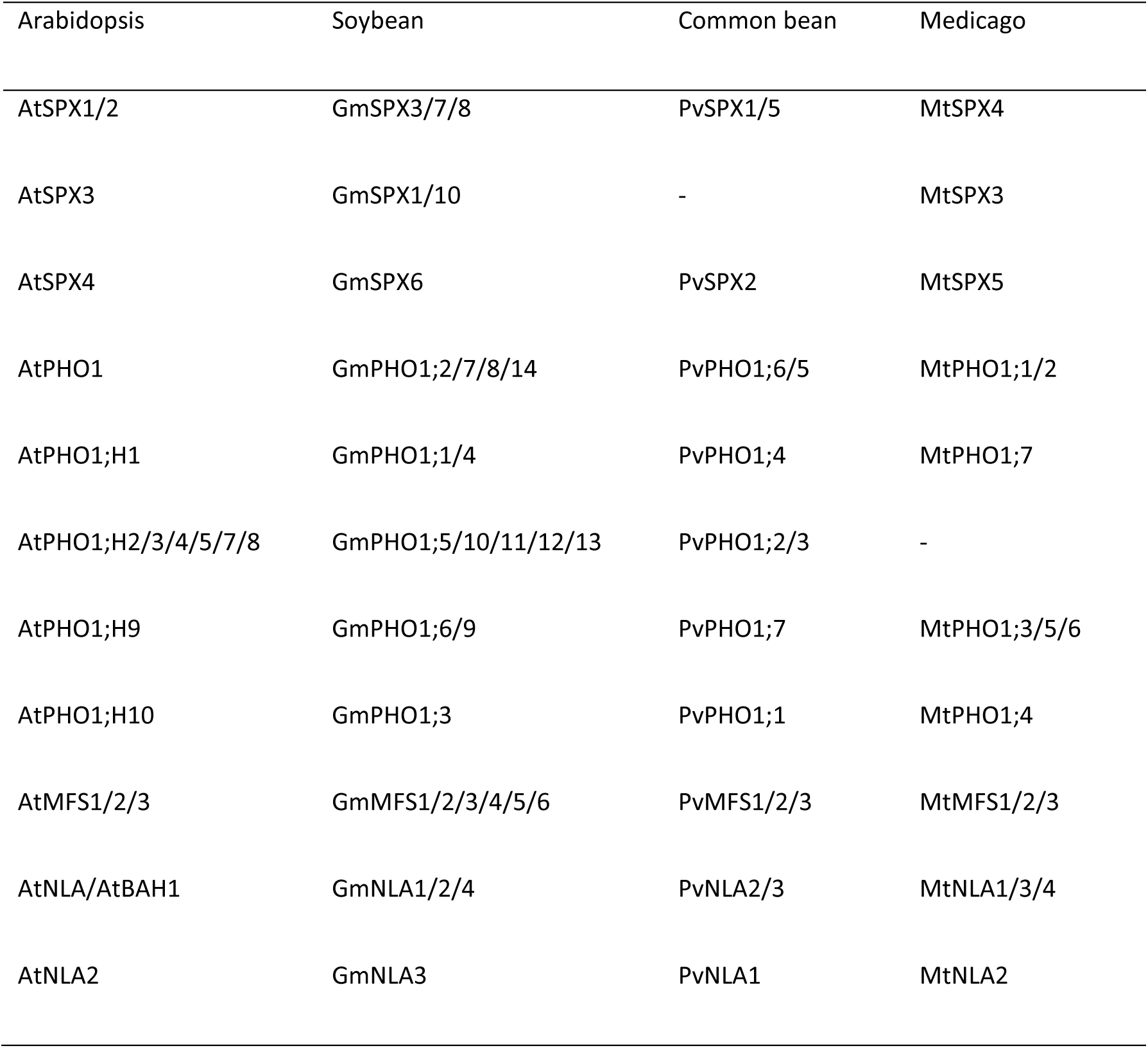
Ortholog genes between legumes and Arabidopsis.

### Expression analysis of SPXs in Arabidopsis and soybean

SPX genes are involved in various physiological process but they are specifically known for their role in phosphate signaling and phosphate homeostasis. To get insight into the potential developmental roles and preferential tissue expression, we analyzed a raw RNA-seq dataset from different developmental stages of different soybean tissues (PRJNA238493). We profiled the *GmSPXs* expression across 17 different samples (Additional file 1: Figure S13). Overall, we observed different expression patterns of *GmSPXs* in various developmental stages of different tissues, indicating a functional divergence in each class of *GmSPXs* [27, 44]. For example, *GmMFS2/5* and *GmPHO1;2/7* showed the same expression in almost all samples but were preferentially expressed in leaf and root, respectively. It can be concluded that they are not involved in the developmental processes. On the other hand, duplicated gene pairs arising from Glycine-specific WGD showed very similar expression patterns across all the samples, especially the *GmMFS2/5* gene pair, but except *GmSPX5/9* and *GmPHO1;5/10* pairs. Taking together, both groups of duplicated genes with the same or different expression pattern showed the evidence of sub-functionalization during the soybean evolution [44].

In order to gain insight how individual SPX genes are regulated by Pi deficiency, we analysed publicly available RNAseq dataset (PRJNA544698) [45] and used DPGP software to cluster genes with similar response patterns. DPGP clustering revealed 6 and 4 clusters for root (Additional file 1: Figure S14) and leaf (Additional file 1: Figure S15), respectively. We designated names for each cluster based on their patterns; up-reg-fast (cluster 3 in root and cluster 1 in leaf), down-reg-fast (cluster 2 in root and cluster 3 in leaf), the lowest-peak-T1 (cluster 6 in root), the lowest-peak-T2 (cluster 5 in root), the highest-peak-T1 (cluster 1 in root), up-reg-slow (cluster 4 in leaf), and the highest-peak-T2 (cluster 4 in root and cluster 2 in leaf). As can be seen in the Table 4, some genes have opposite pattern of regulation in different tissues. To exemplify, *GmSPX1* was placed in down-reg-fast in root and up-reg-fast in leaf, *GmSPX-PHO1;10* is found in the highest-peak-T1 in root and the highest-peak-T2 in the leaf, while *GmSPX6*, *GmSPX-NLA1*, and *GmSPX-NLA3* were in the lowest-peak-T2 cluster in root and the highest-peak-T2 in leaf. The homologs of *AtPHO1* and *AtPHO1;H1(PHO1;2/7/14)* showed an up-reg-fast pattern of cluster 4 in root and the highest-pick-T2 in clusters 2 leaf. Supporting these patterns, He et al. (2013) reported similar expression pattern for these genes, however, there is no clear association between increasing mRNA level of these genes in leaves during phosphate deficiency and growth or shoot Pi content [15]. Overall, for the genes which show tissue-specific expression, we observed different patterns in root and shoot in response to phosphate deficiency.

**Table 4.** Different patterns of clusters in root and leaf in the time series dataset of soybean.

| Patterns | Root clusters | Leaf clusters |
| --- | --- | --- |
| **up-reg-fast** | **Cluster3:** *SPX4, SPX.PHO1;2,*<br>*SPX.PHO1;7, SPX.PHO1;8, SPX.PHO1;14,*<br>*SPX.PHO1;6* | **Cluster1:** *SPX1, SPX.MFS6,*<br>*MFS-NLA2, MFS-NLA4,*<br>*SPX.PHO1;11* |
| **down-reg-fast** | **Cluster2:** *SPX1, SPX10, SPX5, SPX.MFS4,*<br>*SPX.NLA2, SPX.NLA4, SPX.PHO1;12* | **Cluster3:** *SPX2, SPX.MFS2,*<br>*SPX.MFS5, SPX.MFS4* |
| **up-reg-slow** |  | **Cluster4:** *SPX4, SPX5,*<br>*SPX.PHO1;1, SPX.PHO1;4,*<br>*SPX.PHO1;6, SPX.PHO1;9* |
| **highest-peak-T1** | **Cluster1:** *SPX.MFS1, SPX.PHO1;5,*<br>*SPX.PHO1;10, SPX.PHO1;9, SPX.PHO1;1* |  |
| **highest-peak-T2** | **Cluster4:** *SPX.MFS5, SPX.PHO1;4,*<br>*SPX.PHO1;11* | **Cluster2:** *SPX3, SPX6, SPX7,*<br>*SPX8, SPX10, SPX.NLA1,*<br>*SPX.NLA3, SPX.PHO1;2,*<br>*SPX.PHO1;7, SPX.PHO1;14,*<br>*SPX.PHO1;5, SPX.PHO1;10,*<br>*SPX.PHO1;12* |
| **lowest-peak-T1** | **Cluster6:** *SPX3, SPX7, SPX8, SPX.MFS2,*<br>*SPX.MFS6* |  |
| **lowest-peak-T2** | **Cluster5:** *SPX6, SPX.NLA1, SPX.NLA3* |  |

Finally, after investigating developmental and dynamical expression patterns of *GmSPX*, we used another RNA-seq dataset from Arabidopsis and soybean to examine the expression of *SPXs* in three different zones of root [46]. The original data were generated in multiple species, however, we only used RPKM values from Arabidopsis and soybean. A general comparison showed that almost all SPX tended to group species-based rather than orthology-based, except *AtPHO1* and *AtPHO1;H1* which clustered with their orthologs, *GmPHO1;2* and *GmPHO1;7* (Additional file 1: Figure S16). Thus, we can conclude that the tissue-specific genes pose difficulty to identify functional orthologs because of probable tissue inequivalences among species.

## Discussion

The role of SPX domain-containing proteins in Pi homeostasis in Arabidopsis, rice, rapeseed, and wheat and to some extent in soybean and common bean were studied previously [3, 13, 27, 29–31, 35]. While an evolutionary analysis of SPX-EXS [47] and SPX-MFS [24] classes has been reported, as far as we know, the evolution of all classes of SPX gene family from algae to higher plants has not been explored. In addition, despite legume crops requiring a relatively high amount of P, no systematic study of SPX gene family has been reported in legume crops. To close this knowledge gap, we performed a comprehensive search for SPX genes throughout three legume crops, including soybean, *M. truncatula*, and common bean and also algae, liverwort, hornwort, and basal angiosperms to figure out how this gene family originated and expanded during the evolution as well as to identify SPX functional orthologs in legumes.

### Evolutionary conservation and divergence of SPX gene family from algae to legumes

Proteins harboring SPX domain has been reported to form four classes based on their domains. Meanwhile, some other classes have been revealed in the basal plants and algae such as SPX-SLC and SPX-VTC [24]. Here we report other functional protein domains being fused to SPX domains, including EIN3, S6PP, EIN3-S6PP, and Kelch in *S. moellendorffii*, CitMHS in *C. crispus*, Na_sulph_symp in *G. sulphuraria* and *C. reinhardtii*, BET (Bromodomain extra-terminal-transcription regulation) in *P. somniferum*, EXS.rve in *C. braunii*, and Sugar_tr in *M. polymorpha*. Interestingly, some of these new domains have been lost in the land plants and all of them in angiosperms. Domains present in algae before land colonization probably had specific functions that are not required for land plants. For example, SPX-SLC and SPX-VTC were reported in algae that store polyP and are thus lost in plants with Pi vacuole storage, which in turn gained SPX-MFS [24]. Among all assayed species, *S. moellendorffii* showed the most variation of SPX genes, which could be due to its special ability of resurrection. Moreover, unlike other SPX proteins, SPX domain are located at C terminal in S6PP-SPX, EIN3-S6PP_C-SPX, EIN3-SPX, and Kelch-SPX classes. The function of other fusion proteins is unknown so far. Particularly interesting are the fusions of SPX with EIN3 domains, because in Arabidopsis EIN3 is directly involved in regulation of phosphate homeostasis through binding to promoter of *PHR1* [48]. The SPX domain would then add another level of control for this interaction and allow the reciprocal regulation of ethylene signaling by phosphate. Similarly, Kelch domains are often found in regulatory proteins, for example fused to F-Box proteins [49], hence, again, the fusion with SPX may connect multiple regulatory circuits. If the SPX domain enables the activities of the additional domains to be modulated by phosphate (or InsPP), this offers an intriguing opportunity for using these domains in synthetic biology approaches to make various cellular processes controlled by phosphate. These hypotheses, however, have to be verified. On the other hand, RING and MFS classes have gradually appeared in the later-diverging plants. MFS and then RING class have the least fluctuations from 1 to 6 genes. In contrast, EXS class had high variation of gene numbers in each species and also the highest number of identified genes in comparison with the other classes. Also, presence of this domain in whole Eukarya except algae, suggest that it has been lost in some algae.

The number of whole-genome duplications is correlated with gene family size [47, 50], which is consistent with our results, since *P. somniferum* and *G. max* with two WGD events had the largest sizes of SPX family [51, 52]. The expansion of SPX family in these two plants is mostly affected by WGD duplication type, while segmental/local duplication type was the main contributor of expansion in *S. moellendorffii*, the species with third greatest SPX family, which might explain its unique classes. Algae possess 2 to 5 SPX gene family members. The expansion in *P. patens* (22 members), could suggest that duplications took place after plant terrestrialization as the SPX proteins became more important [53].

The phylogenetic analysis brought some unexpected findings. First, it showed three clades for 4 subfamilies; SPX and EXS in two different clades, but MFS and RING classes diverged from the same ancestor. Second, SPXs from algae did not group with other species in any clade, except of SPX-I. It can be concluded that genes in the SPX-I sub-clade are the most ancient genes in angiosperms that were diverged from the same ancestor with green algae. Hence, *AtSPX4*, *GmSPX6*, *MtSPX5*, *PvSPX2*, and *OsSPX4* probably have the same function with their ancestral orthologs in the green algae, but the genes in two other sub-clades, SPX-II and SPX-III have evolved after divergence of streptophytes and chlorophytes and might have acquired additional functions. AtSPX4 and OsSPX4 have indeed the same function and mechanism in regulation of PSI, as in presence of phosphate both proteins interact in the cytosol with the corresponding key regulators AtPHR1 and OsPHR2, and prevent them from translocating to nucleus [11, 54]. During P deficiency they are rapidly degraded, releasing thus the PHR factors to induce transcription of PSI genes. The two proteins however, also differ, as while OsSPX4 integrates nitrate and phosphate signaling, AtSPX4 does not seem to have this function, but on the other hand integrates phosphate signaling and anthocyanin biosynthesis [11, 55].

The MFS class as the most recently diverged class of SPX proteins was divided into two sub-clades, with MFS-I specifically containing monocots, suggesting that MFS genes in monocots diversified differently in comparison with basal angiosperms and eudicots. This may be due to different Pi storage between monocots and eudicots [6, 56, 57]. While monocots store P preferentially in the roots and their leaves have the highest P concentration in the mesophyll cells, eudicots store much more P in the leaves with the highest concentration in the epidermis [56]. It is thus possible that the different cellular localization drove a different evolution of SPX-MFS genes between monocots and dicots. Modern RING class genes have evolved two times, RING-I clade arose from a duplication of the common ancestor of mosses and angiosperms and RING-II arose from duplication of the common ancestor of lycophytes, liverwort and angiosperms. In the EXS class, EXS-III clade did not contain any orthologs from monocots but interestingly, many *AtPHO1* genes such as *AtPHO1;H2/3/4/5/6/7/8* grouped specifically with the genes from *Brassica napus*, suggesting special functions in Brassicaceae. Only *AtPHO1;H9* and *AtPHO1;H10* had two and one orthologs in the legumes, respectively.

Based on collinearity analyses, species with more WGD events showed more inter-species collinearity, but *S. moellendorffii* with locally expanded SPX and rice with mostly dispersed expanded SPX just showed intra-genome collinearity. Low collinear relationship between rice and eudicots was reported previously [58] and explained by longer evolutionary distance and more genome rearrangements [59] as well as the erosion of macrosynteny between monocots and dicots [60]. Our results are consistent with the monocot paleopolyploidy after their divergence from eudicots [58]. Having collinear relationship can arise from paleopolyploidy in the common ancestor, but *S. moellendorffii* has no evidence for WGD events and its intra-genome collinear blocks arose from segmental/local duplication [37].

### Functional characterization of SPXs in legumes

Due to the functional conservation of proteins across species, determination of orthologous relationships can provide useful insights about the biological role of these proteins [61]. As plants have undergone various duplication events and had different evolutionary trajectories, relating same functions to the orthologs are difficult, especially there are one-to-many or many-to-many orthologous relationships [32]. Therefore, two different methods, phylogenetic inference of orthologs from protein sequences and expressolog identification, were conducted for prediction of functional orthologs of SPXs. This was necessary because, firstly, there are complex orthology relationships among some SPX genes that prevented Orthofinder to detect the exact functional orthologs and, secondly, some SPX genes show tissue-expression pattern that can pose problem to identify expressologs, due to difficulties in assignment of tissue equivalencies between legumes and Arabidopsis. In the dynamic *GmSPX* expression patterns, we observed tissue-specificity for most of *GmSPX*s except for homologs of *AtPHO1* and *AtPHO1;H1*. Taking together, we could assign functions of *AtSPX4*, *AtPHO1;H10* and *AtNLA2* to their predicted orthologs from Orthofinder and *AtPHO1* and *AtPHO1;H1* to their orthologs from expressolog identification results. To examine this conclusion, we analyzed two different datasets of soybean to profile *GmSPX*s expression in different tissues and developmental stages as well as their dynamic expression responses to Pi deficiency in leaf and root. Overall, we found that almost all *GmSPX*s except *GmPHO1;2/7* and *GmMFS2* have different expression patterns across the developmental samples as well as in root and leaf responses to the dynamic Pi deficiency. In summary, these transcriptome analyses highlighted that GmSPX genes might be involved in different developmental processes and stresses beyond phosphate starvation response. It is probable that new or sub-functionalization in soybean and generally in legumes took place with the new functions of SPX proteins waiting to be discovered. Our analyses lay a solid foundation for the future functional studies of SPX proteins from algae to legumes.

### Conclusion

In conclusion, we comprehensively analyzed SPX gene family evolution and dissected how different protein motifs and Cis-acting elements evolved, as well as identified expansion patterns, and collinear gene blocks during evolution from algae to angiosperms. Afterwards, focusing on legumes, we tried to model evolutionary history of SPXs in soybean and identify functional orthologs. We could predict the putative SPX proteins involved in long-distance Pi transportation in soybean and Medicago. Our study not only provides a global view of the evolution and expansion of *SPX* gene family in important species but also provides the first step for more detailed investigations of the functions of individual *SPXs* in legumes.

## Material and methods

### Bioinformatic identification of SPX proteins

In order to identify SPX domain-containing proteins in our species; legume crops (soybean – *Glycine max*, alfalfa – *Medicago truncatula*, and common bean – *Phaseolus vulgaris*), mosses (*Physcomitrella patens*), liverwort (*Marchantia polymorpha*), Rhodophytes (*Cyanidioschyzon merolae*, *Galdieria sulphuraria*, and *Chondrus crispus*), chlorophytes (*Chlamydomonas reinhardtii* and *Ostreococcus lucimarinus*) , charophytes (*Chara braunii*), basal angiosperms (*Papaver somniferum*, *Amborella trichopoda*, and *Nymphaea colorata*), and lycophytes (*Selaginella moellendorffii*), full-length protein sequences of AtSPXs were used for BLASTP searches across proteomes of the above mentioned species. After removing redundant sequences, the SPX proteins obtained through BLASTP search were investigated for the presence of additional domains along with SPX domain using SMART [62], Pfam [63], Conserved Domain Database (CDD) [64], and PROSITE [65] databases.

The sequences of identified SPX proteins in the three legume crops were analyzed for their physiochemical properties; including isoelectric point (pI), molecular weight (Mw), instability index (II), grand average of hydropathicity (GRAVY), and aliphatic index (AI) using ProtParam tool of ExPASy website (https://web.expasy.org/protparam/). Subcellular location prediction was conducted using Wolf Psort [66].

### Phylogeny analysis and identification of conserved motifs

The amino acid sequences of identified SPX proteins in our surveyed species and Arabidopsis as reviewed in [6], rice [6], wheat [3], and *Brassica napus* [27] were downloaded from EnsemblPlants (https://plants.ensembl.org/index.html). Three sequences to be used as outgroup, XPR1 from human and mouse, and SYG1 from *C. elegans,* were downloaded from NCBI database (https://www.ncbi.nlm.nih.gov/). Multiple sequence alignment of these full-length sequences was performed by ClustalX (ver. 2.1; http://www.clustal.org/). Then, we used Maximum Likelihood method and JTT matrix-based model in MEGA 7 software to build a phylogenetic tree from the sequence alignment using following parameters: p-distance model, partial deletion and 1000 bootstraps. To predict conserved motifs of SPX proteins across all species, as well as Arabidopsis and rice, MEME (http://meme-suite.org/tools/meme) tool with the maximum number of motifs 20 was used. Logo sequences of conserved motifs were obtained by Weblogo 3 (http://weblogo.threeplusone.com/).

### Collinearity analysis and gene expansion pattern of SPX from algae to eudicots

In order to get insight about how collinear blocks have been conserved during the evolution, we performed collinearity analysis three times with different species; 1. Among three legume crops, Arabidopsis, rice, *P. somniferum*, and *N. colorota*, 2. Among *S. moellendorffii* , *P. patens*, *N*. *colorata*, and *A. trichopoda*, and 3. Among three legume crops using MCScanX toolkit [67] to get collinear gene blocks and also duplication types by duplicate_gene_classifier program. To visualize the collinear blocks among the first and third runs, tbtools was used [68]. Because of non-chromosomal reference genomes in *P. patens* and *S. moellendorffii* we just retrieved their collinear gene blocks without visualization.

### Selective pressure and evolutionary models of SPX genes in the legume crops

Duplication blocks between each two species of soybean, common bean and *M. truncatula* were retrieved from the Plant Genome Duplication Database (PGDD, http://chibba.agtec.uga.edu/duplication/). SPX gene blocks were manually extracted and used for further analyses. The selective pressure on duplicated genes were estimated by retrieving synonymous (Ks) and non-synonymous (Ka) per site between the duplicated gene-pairs using from PGDD database. The Ka/Ks ratio was assessed to determine the molecular evolutionary rates of each gene pair. Generally, the Ka/Ks<1 indicates purifying selection, Ka/Ks>1 indicates positive selection, and Ka/Ks=1 indicates neutral selection. The divergence time of the duplication blocks was evaluated to investigate the evolution of GmSPX genes. If the Ks > 1.5, the divergence time is after the Gamma whole-genome triplication (WGT); if the Ks < 0.3, the divergence time is after the Glycine whole-genome duplication (WGD) event; and when the Ks is between 0.3 and 1.5, the divergence time is after legume WGD event but before the Glycine WGD event [69, 70].

### Identification of Cis-acting-elements in the promoters of SPX gene family

For finding evolutionary pattern of Cis-acting-elements from algae to eudicots, 1500 bp upstream from the start codon of SPX genes in all assayed species and Arabidopsis were downloaded from the EnsemblPlants and analyzed using the PlantCARE database (http://bioinformatics.psb.ugent.be/webtools/plantcare/html/). Afterwards, SPX genes were clustered with hierarchical clustering on principal components (HCPC) method by FactMineR package. All detected cis-acting elements were merged into one matrix with 1 and 0 values for present or absent elements in each promoter, respectively.

### Prediction of functional orthologs of AtSPXs across legumes

To identify functional orthologs in the three legumes, we used OrthFinder to compare SPX genes among 7 species (rice, wheat, rapeseed, Arabidopsis, *M. truncatula*, soybean, and common bean), resulting in orthogroups and orthologs based on sequence similarities [71]. Then, to overcome the weakness of sequence-based ortholog identification for one-to-many and many-to-many orthologs, expressolog identification among Arabidopsis, soybean, and Medicago (http://bar.utoronto.ca/expressolog_treeviewer/cgi-bin/expressolog_treeviewer.cgi), was used.

### Expression analysis of SPX genes

Three different expression analyses were performed as follows:

1. To compare tissue and developmental expression pattern of *GmSPX*s, RNA-seq data of 17 samples from different tissues (flower, root, shoot meristem, seed, and leaves) in five developmental stages (germination, trefoil, flowering, seed development, and plant senescence) (PRJNA238493) [72] were analyzed. The gene expression profiles were visualized by heatmap using R package pheatmap (https://www.r-project.org/).
2. To visualize changes in *GmSPX* gene expression in response to P deficiency we used publicly available dataset (PRJNA544698) [45]. The data were reanalyzed and TPM (Transcript Per Million) values were calculated from samples over different time points of Pi deficiency, including early stress (T, 24 h), recovery (TC, 24 h deficiency, 48 h resupply), and repeated stress (TCT, additional 24 h deficiency) in root and leaf tissues. The data were clustered using the Dirichlet process with Gaussian process mixture model (DPGP) [73].
3. To assess if the predicted functional orthologs in Arabidopsis and soybean show the same expression in different root development zones, including meristemic zone (MZ), elongation zone (EZ), and differentiation zone (DZ) data from [46] have been used. RPKM values for the *SPX*s were collected (GSE64665), and log2 (RPKM + 1) was used to construct correlation heatmap using the pheatmap package (https://www.r-project.org/).

## Declarations

### Ethics approval and consent to participate

Not applicable

### Consent for publication

Not applicable

### Availability of data and materials

The datasets generated and/or analyzed during the current study are included in the supplemental material.

### Authors’ contributions

MNC, AN, EE, and SK designed the study. MNC performed the analyses. Assisted with interpretation. MNC wrote the manuscript. All authors reviewed the manuscript. The authors read and approved the final manuscript.

## Acknowledgements

We acknowledge the Ministry of Science, Research and Technology of Iran and the University of Shiraz for the exchange scholarship to University of Cologne

## Abbreviations

PHR: Phosphate starvation Response
PP-InsPs: inositol pyrophosphates
MFS: Major Facilitator Superfamily
RING: Really Interesting New Gene
NLA: Nitrogen Limitation Adaptation
PSI: Pi starvation-induced
Ein3: Ethylene intensive 3
VTC: vacuolar transporter chaperone
CitMHS: Citrate transporter
Na_sulph_symp: sodium sulphate symporter
S6PP_C: Sucrose-6F-phosphate phosphohydrolase C-terminal
Kelch: Galactose oxidase
rve: Integrase core domain
GRAVY: grand average of hydropathicity
LSC: Lysine Surface Cluster
Gm: Glycine max
Mt: Medicago truncatula
Pv: Phaseolus vulgaris
Ps: Papaver sumniferum
Nc: N. colorota
At: Arabidopsis thaliana
Os: Oryza sativa
CRE: cis-acting elements
MeJA: methyl jasmonate
DPGP: Dirichlet process with Gaussian process mixture model
HCPC: hierarchical clustering on principal components

---

**Figure S1.**
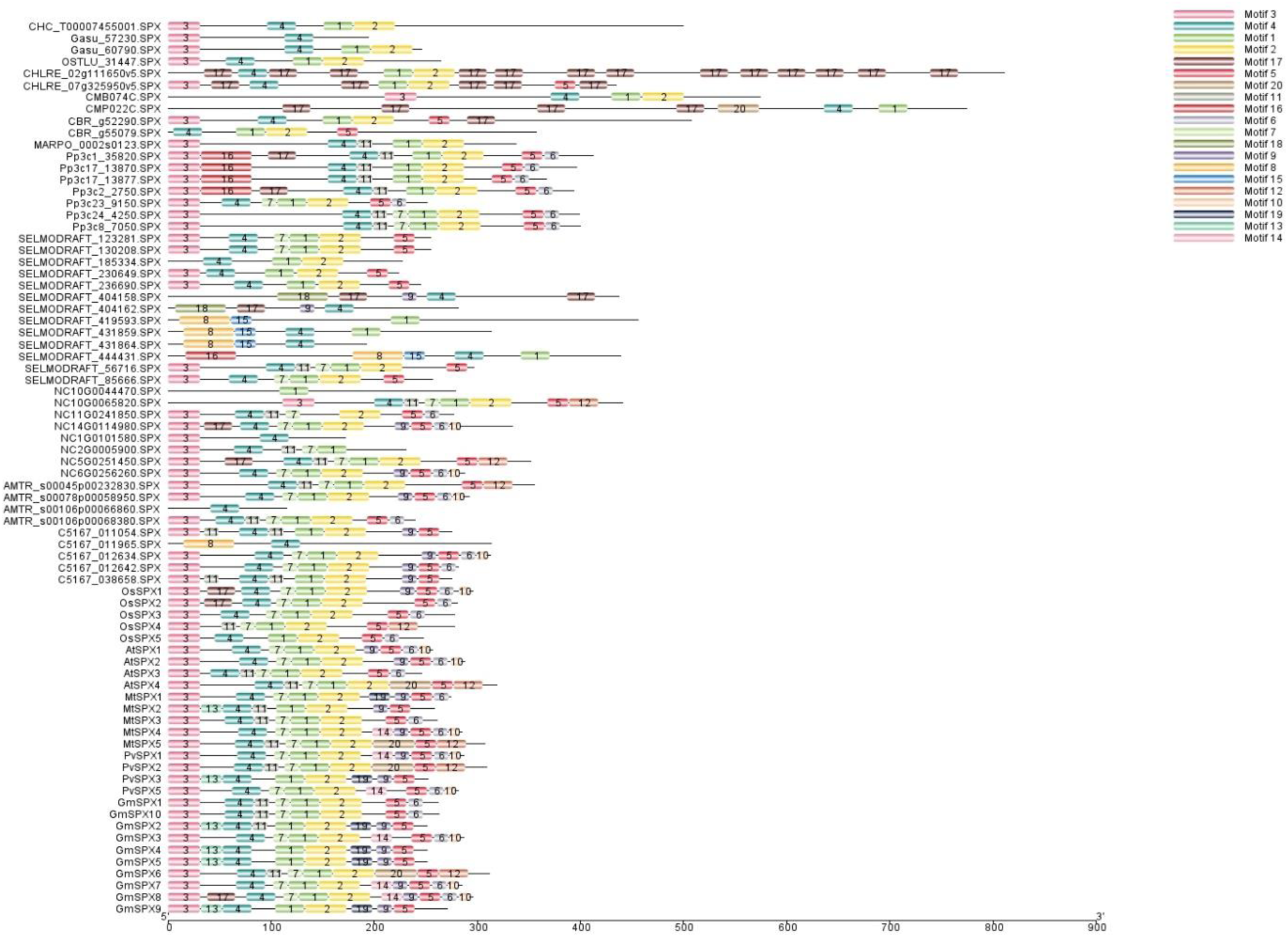
Motif loss and gain in SPX class genes during the evolution from algae to current Angiosperms

**Figure S2.**
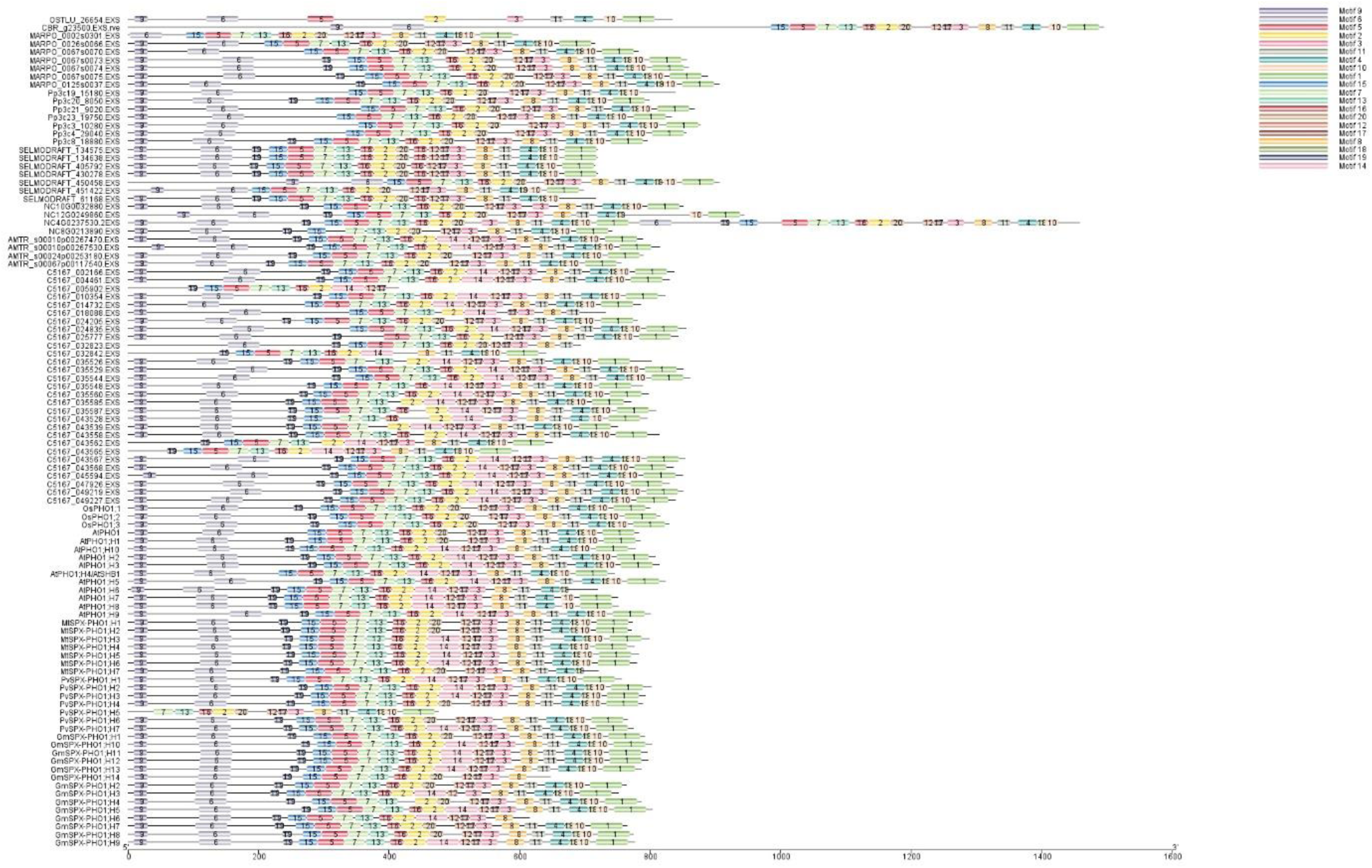
Motif loss and gain in SPX-EXS class genes during the evolution from algae to current Angiosperms

**Figure S3.**
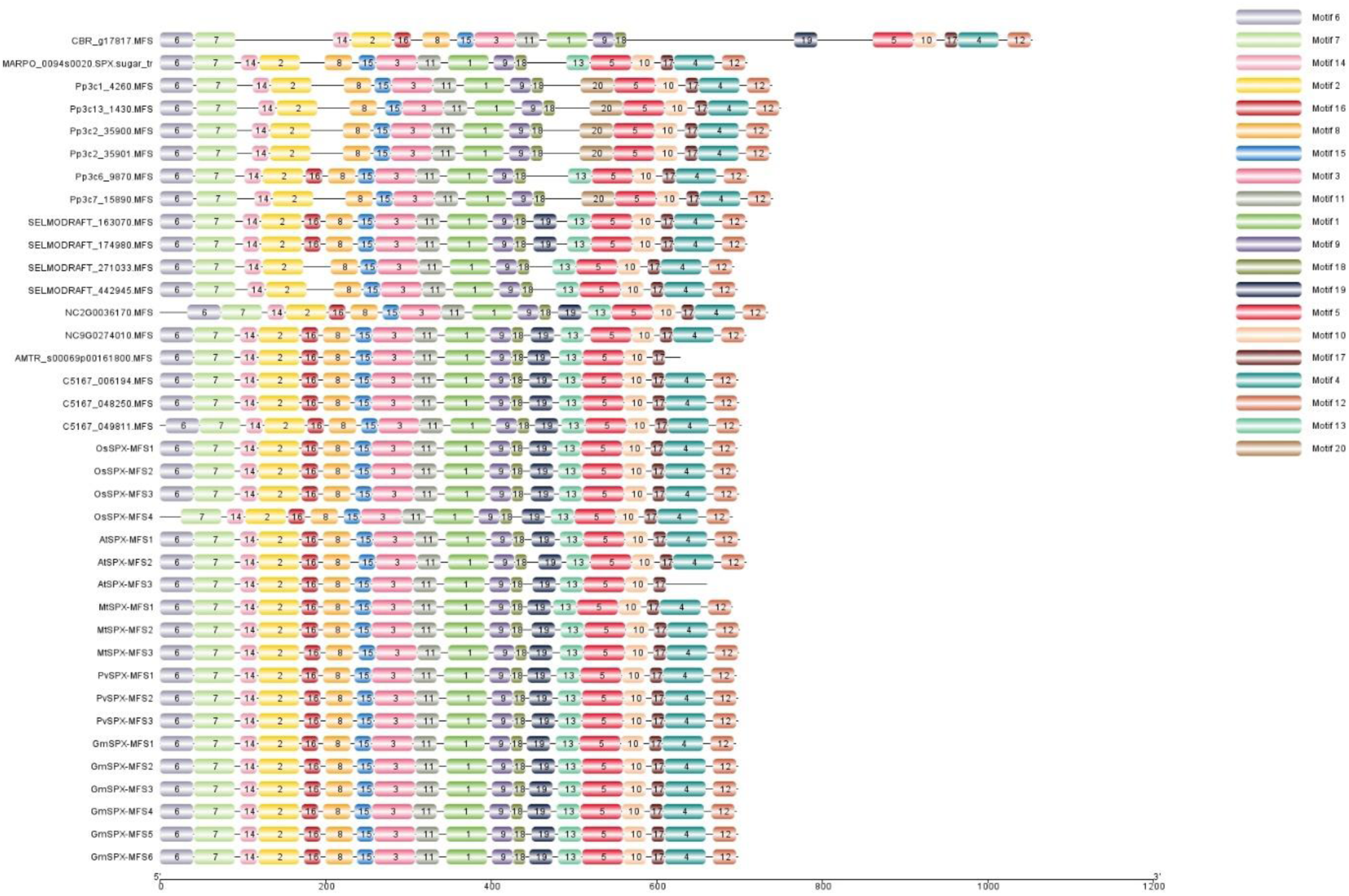
Motif loss and gain in SPX-MFS class genes during the evolution from algae to current Angiosperms

**Figure S4.**
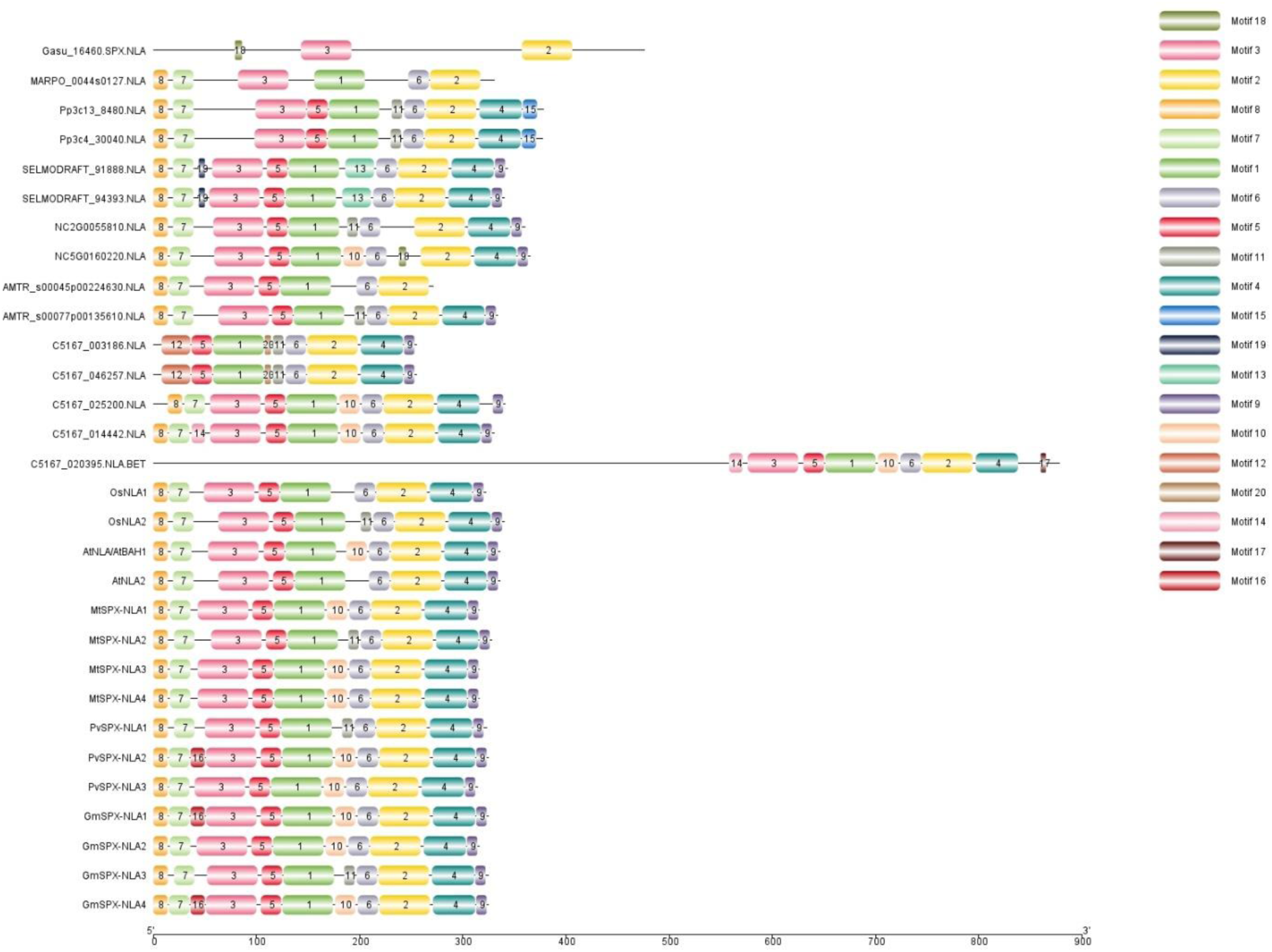
Motif loss and gain in SPX-RING class genes during the evolution from algae to current Angiosperms

**Figure S5.**
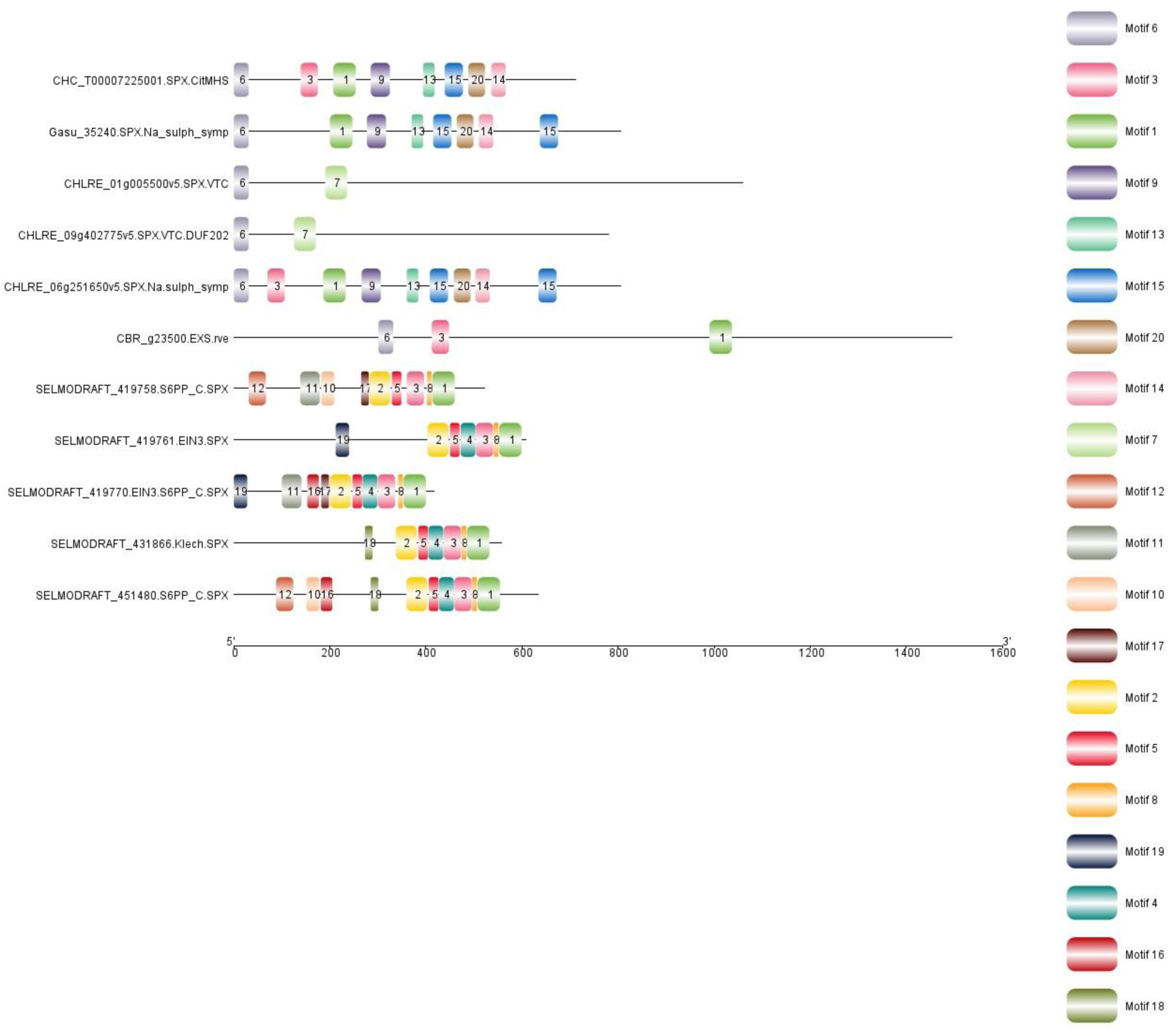
Motifs specifically-found in the new classes of SPX proteins in basal plants

**Figure S6.**
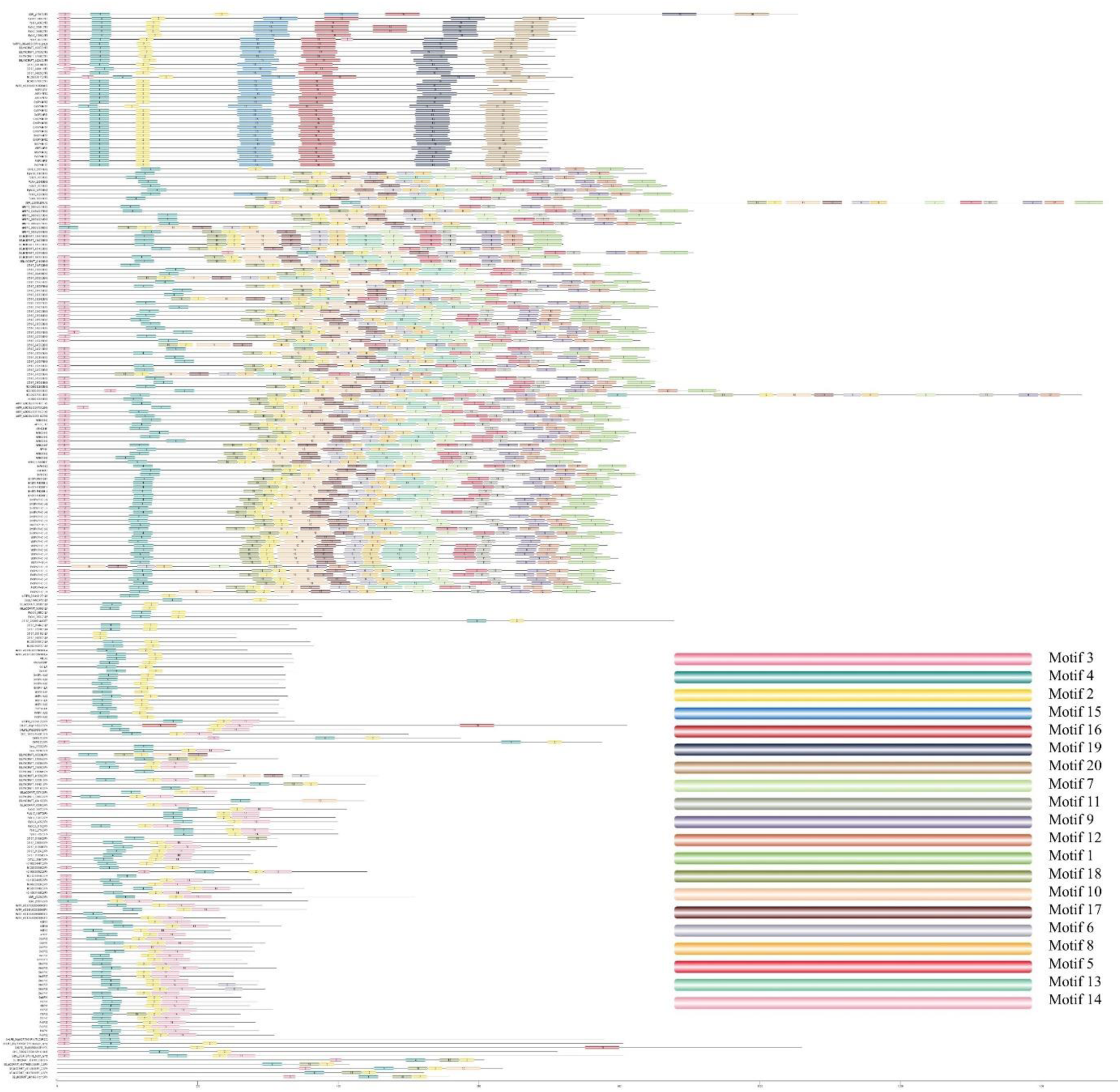
Motif loss and gain of all SPX proteins during the evolution from algae to current Angiosperms

**Figure S7.**
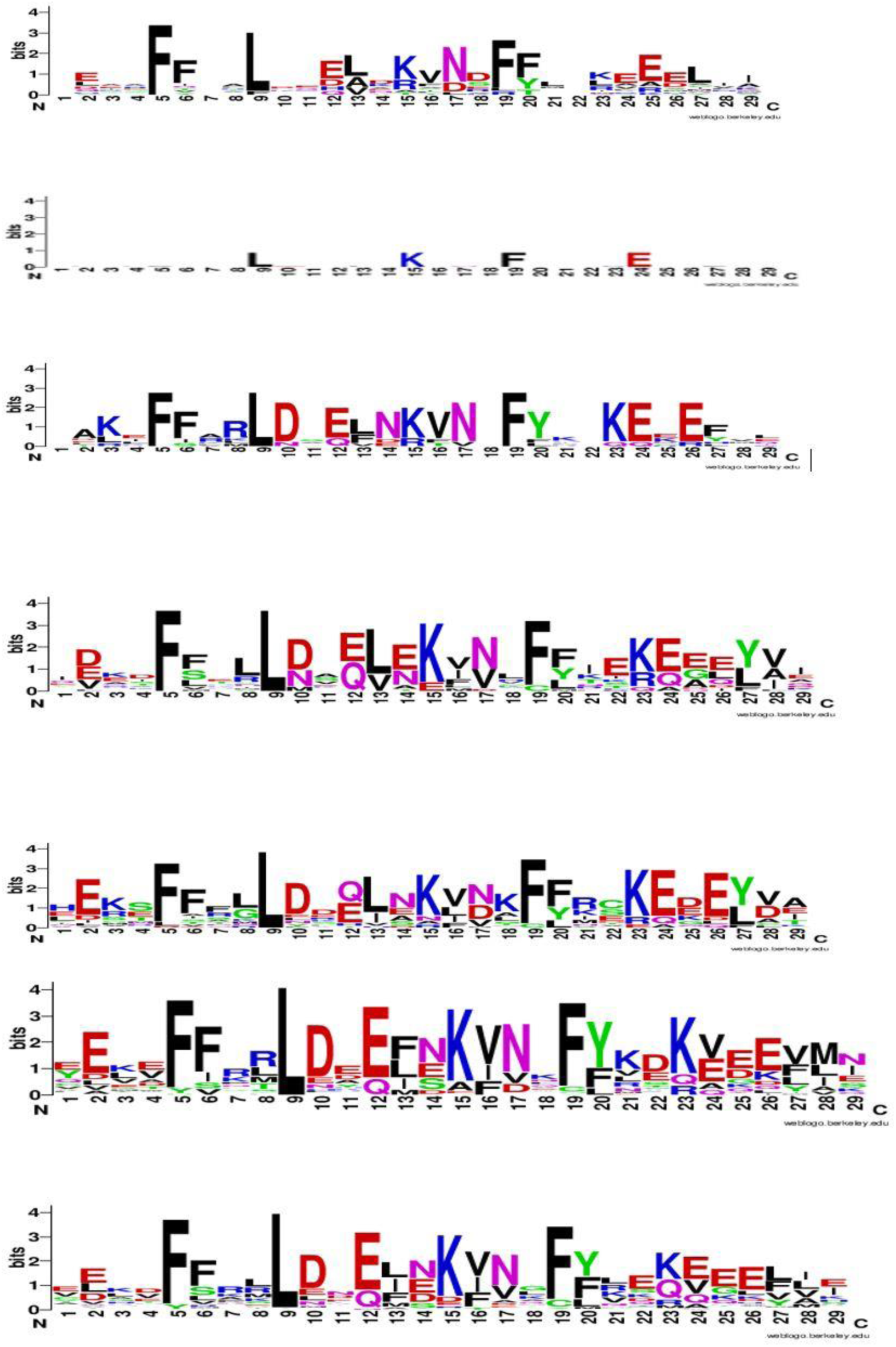
Consensus sequences of motif 4 in SPX domain conserved in whole SPX proteins; in different phyla. Order of phyla from up to down: algae (*C.reinhardttii*, *O. lucimarinus*, *G. sulfuraria*, C*. crispus*, *C. merolae*), charophytes (*C. braunii*), liverwort (*M. polymorpha*), bryophytes (*P. patens*), lycophytes (*S. moellendorffii*), basal angiosperms (*A. thricopoda*, *P. sumniferum*, *N. colorata*), and current angiosperm (Arabidopsis, rice, soybean, common bean, alfalfa).

**Figure S8.**
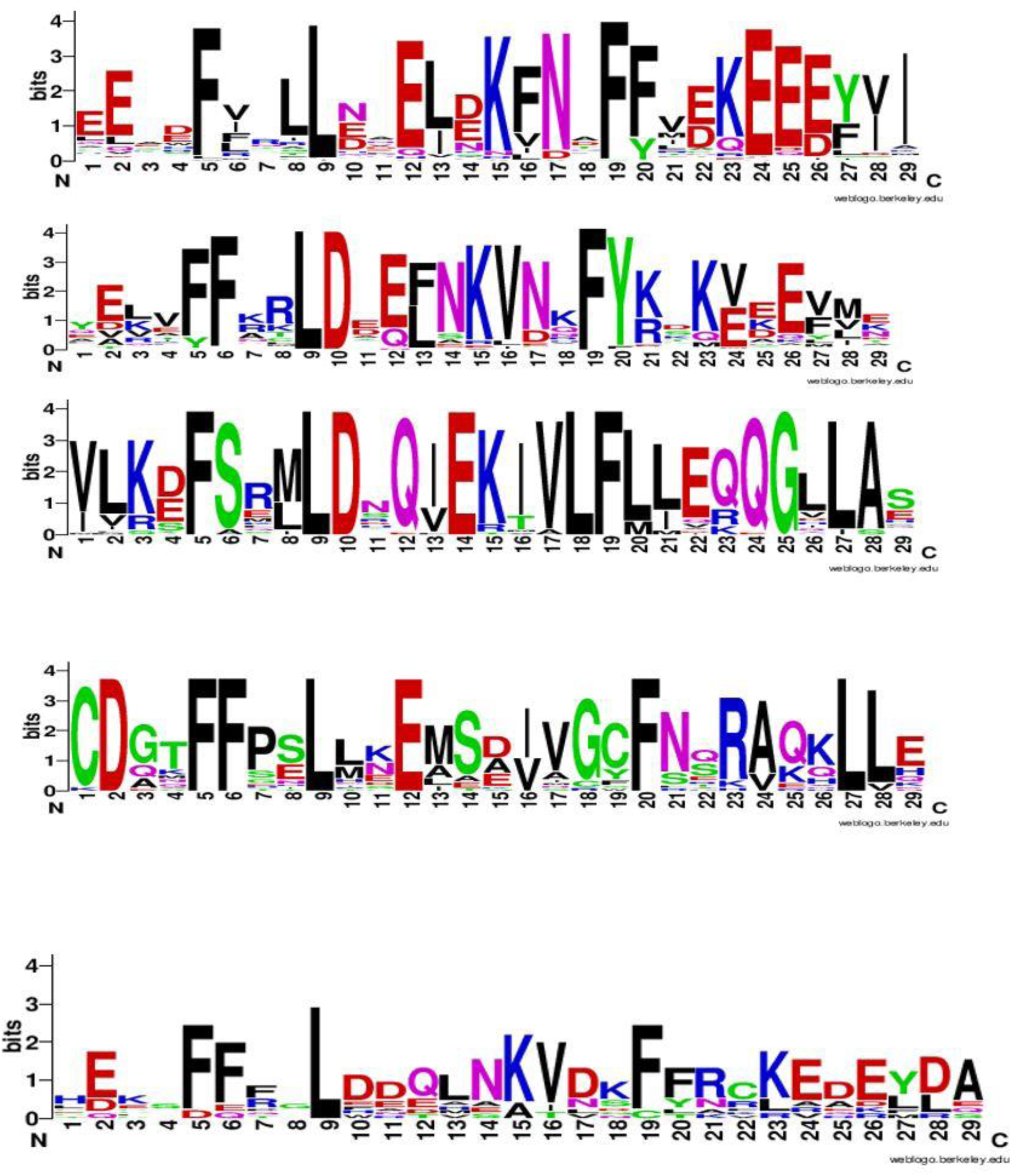
Consensus sequences of motif 4 in SPX domain conserved in whole SPX proteins; in different classes. Order of different classes from up to down: SPX, EXS, MFS, RING, new identified classes.

**Figure S9.**
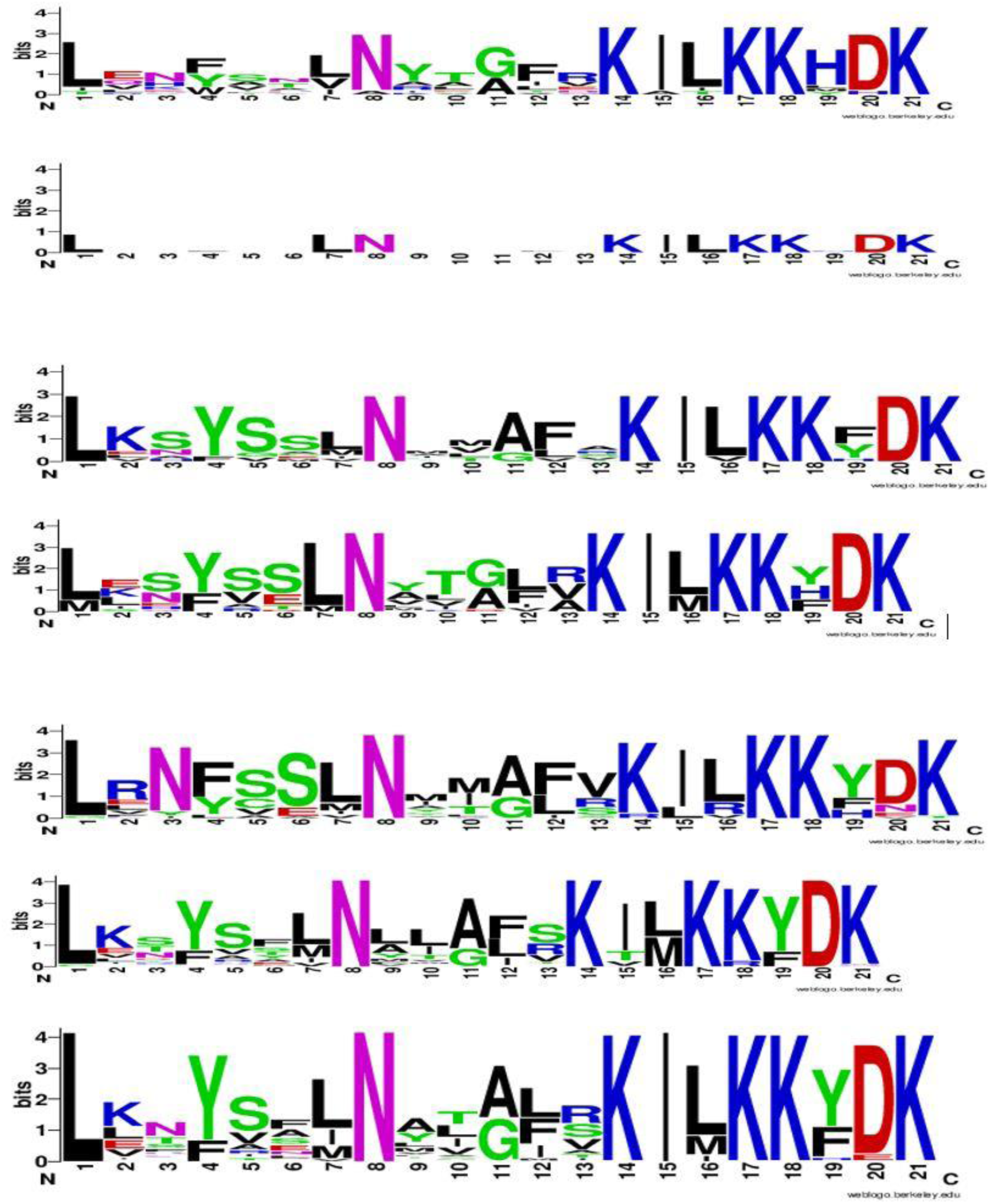
Consensus sequences of motif 2 in SPX domain conserved in whole SPX proteins; in different phyla. Order of phyla from up to down: algae (*C.reinhardttii*, *O. lucimarinus*, *G. sulfuraria*, C*. crispus*, *C. merolae*), charophytes (*C. braunii*), liverwort (*M. polymorpha*), bryophytes (*P. patens*), lycophytes (*S. moellendorffii*), basal angiosperms (*A. thricopoda*, *P. sumniferum*, *N. colorata*), and current angiosperm (Arabidopsis, rice, soybean, common bean, alfalfa).

**Figure S10.**
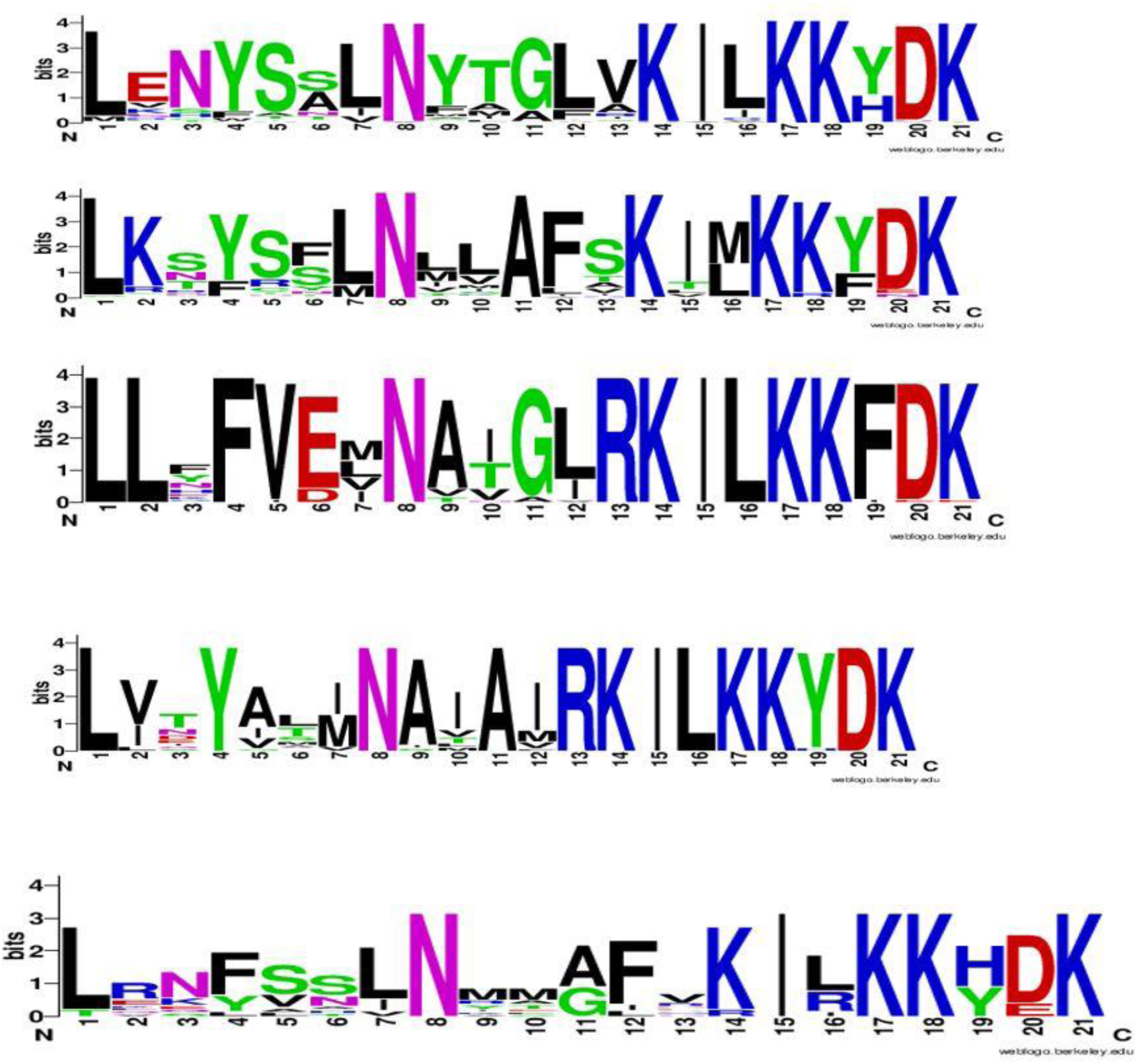
Consensus sequences of motif 2 in SPX domain conserved in whole SPX proteins; in different classes. Order of different classes from up to down: SPX, EXS, MFS, RING, new identified classes.

**Figure S11.**
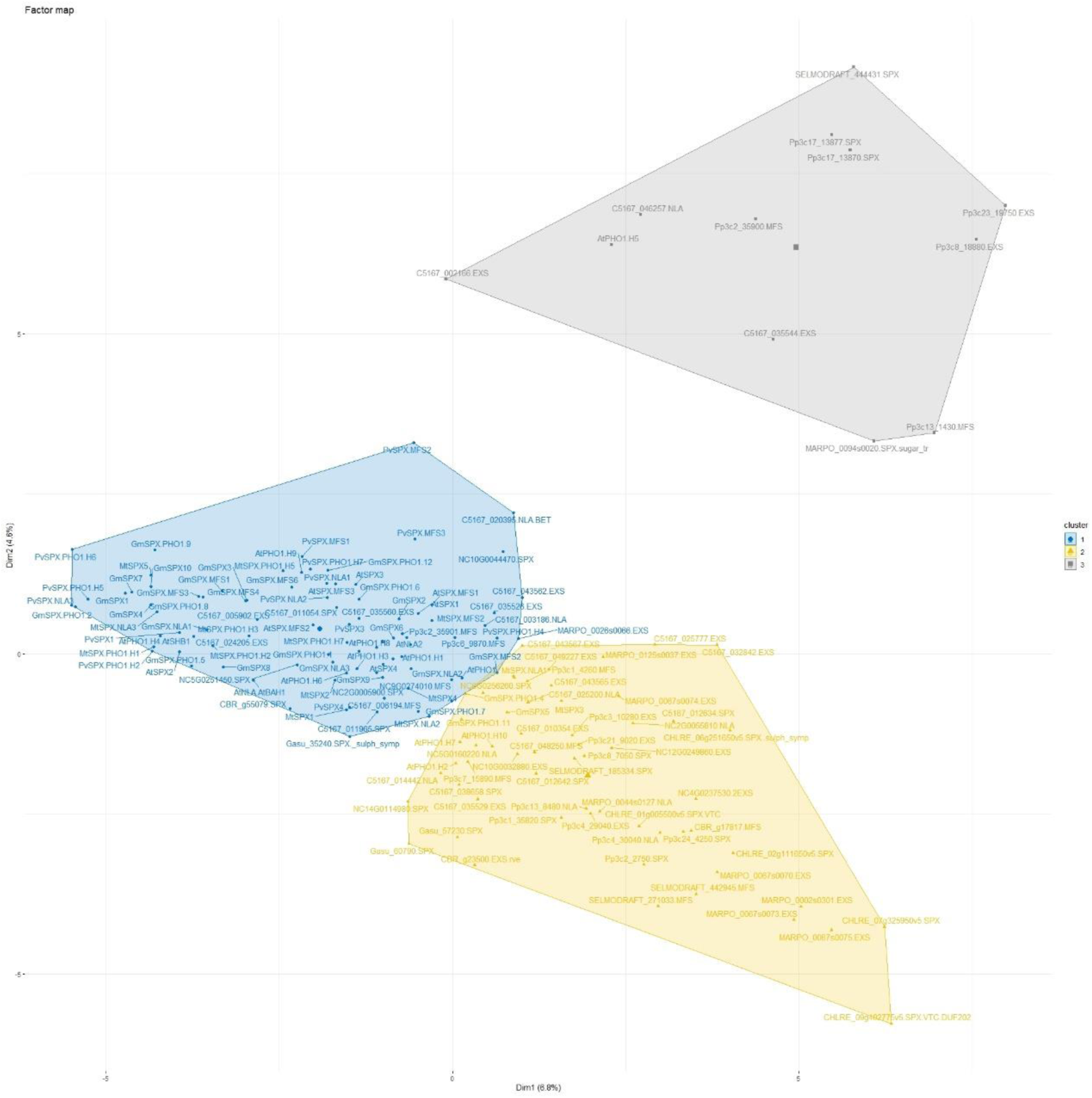
Hierarchical Clustering on Principal Components (HCPC) of SPXs in the lower plants and current Angiosperms based on presence or absence of Cis-acting elements in their promoters.

**Figure S12.**
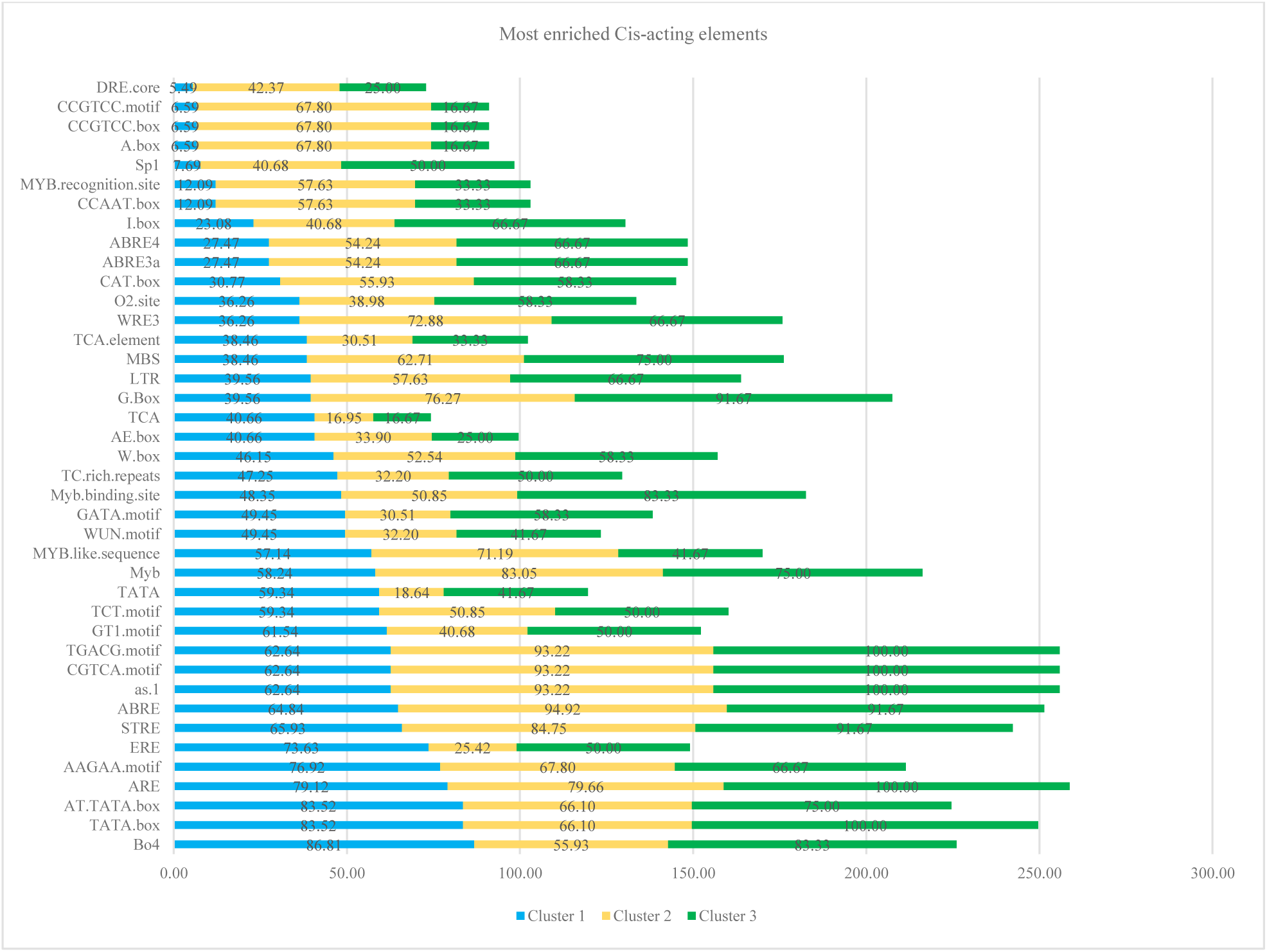
Production of genes in each cluster containing the most frequent Cis-acting elements. The clusters were shown in different colors: cluster 1= blue, cluster 2= yellow, and cluster 3= green.

**Figure S13.**
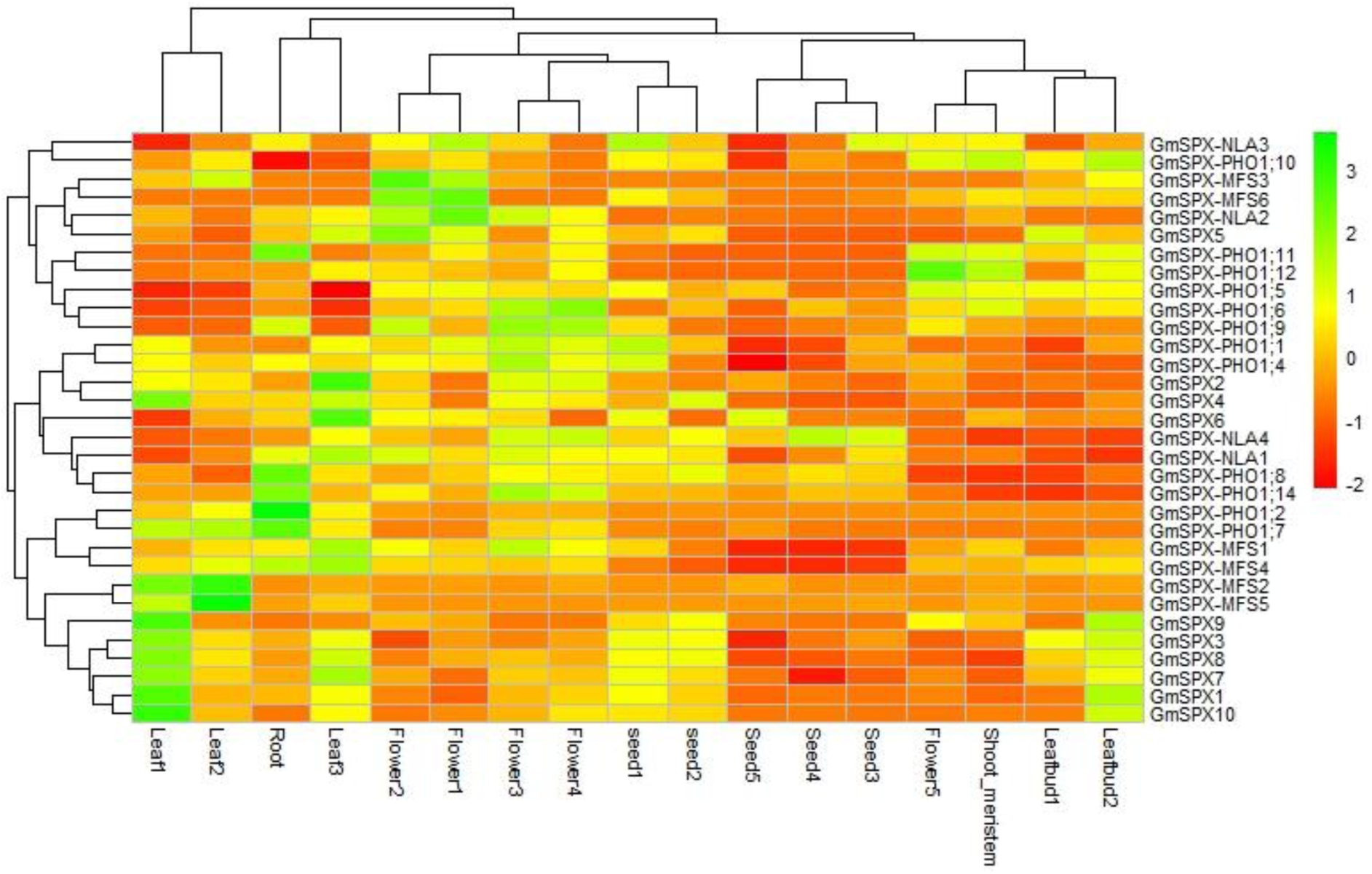
Expression levels of *GmSPXs* in the different developmental stages of different tissues. Using data from PRJNA238493 bioproject.

**Figure S14.**
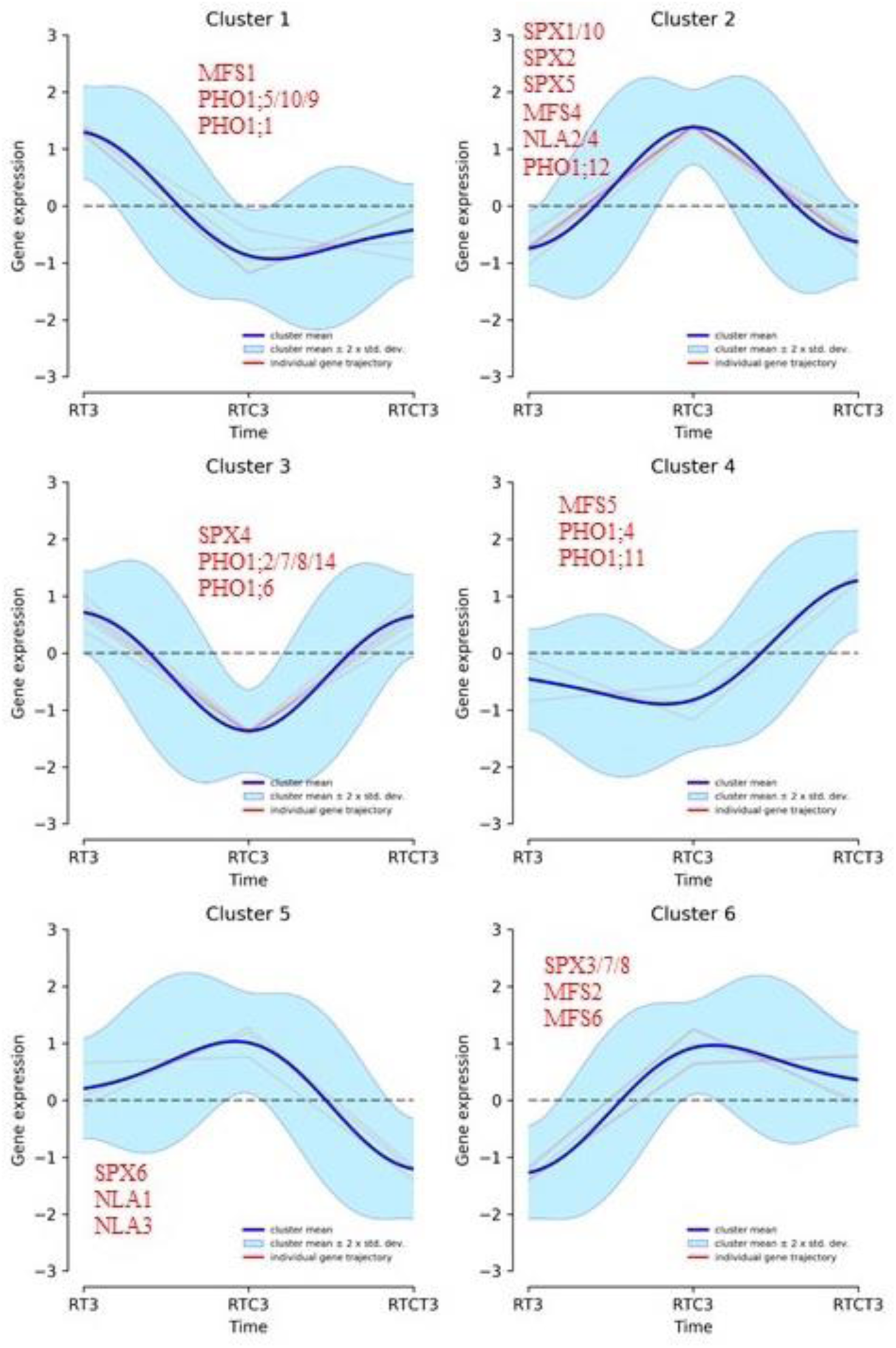
Regulation of SPX genes by phosphate starvation in the roots. DPGP analysis was performed for expression pattern of GmSPXs in roots during three time-points; RT= P deficiency, RTC= P deficiency and recovery, and RTCT = P deficiency, recovery, and second P deficiency. Shown are clustered trajectories of GmSPX genes. The cluster means are in blue, the individual SPX genes are shown in red. Using data from PRJNA544698 bioproject.

**Figure S15.**
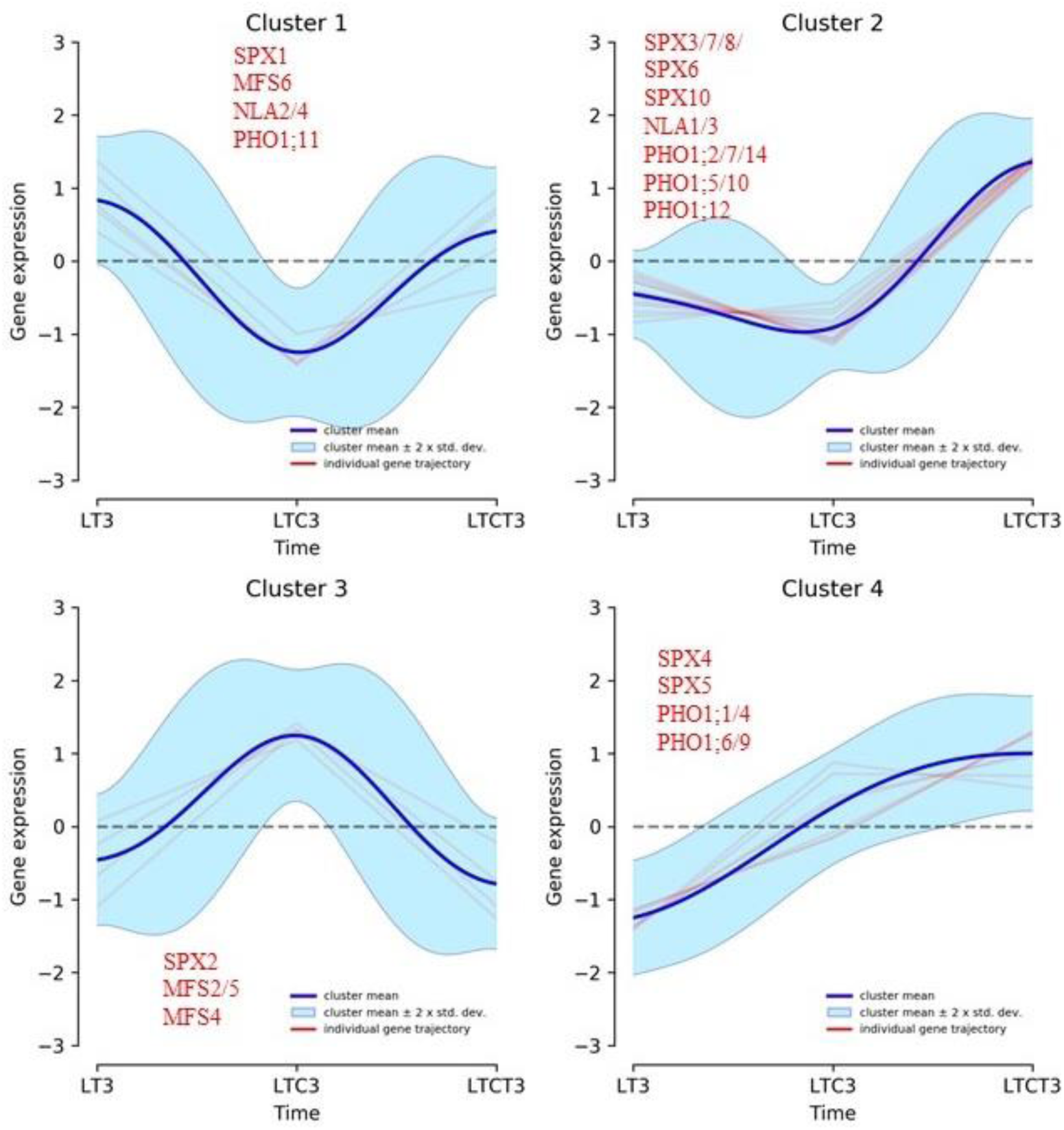
Regulation of SPX genes by phosphate starvation in the leaves. DPGP analysis was performed for expression pattern of GmSPXs in leaves during three time-points; RT= P deficiency, RTC= P deficiency and recovery, and RTCT = P deficiency, recovery, and second P deficiency. Shown are clustered trajectories of GmSPX genes. The cluster means are in blue, the individual SPX genes are shown in red. Using data from PRJNA544698 bioproject.

**Figure S16.**
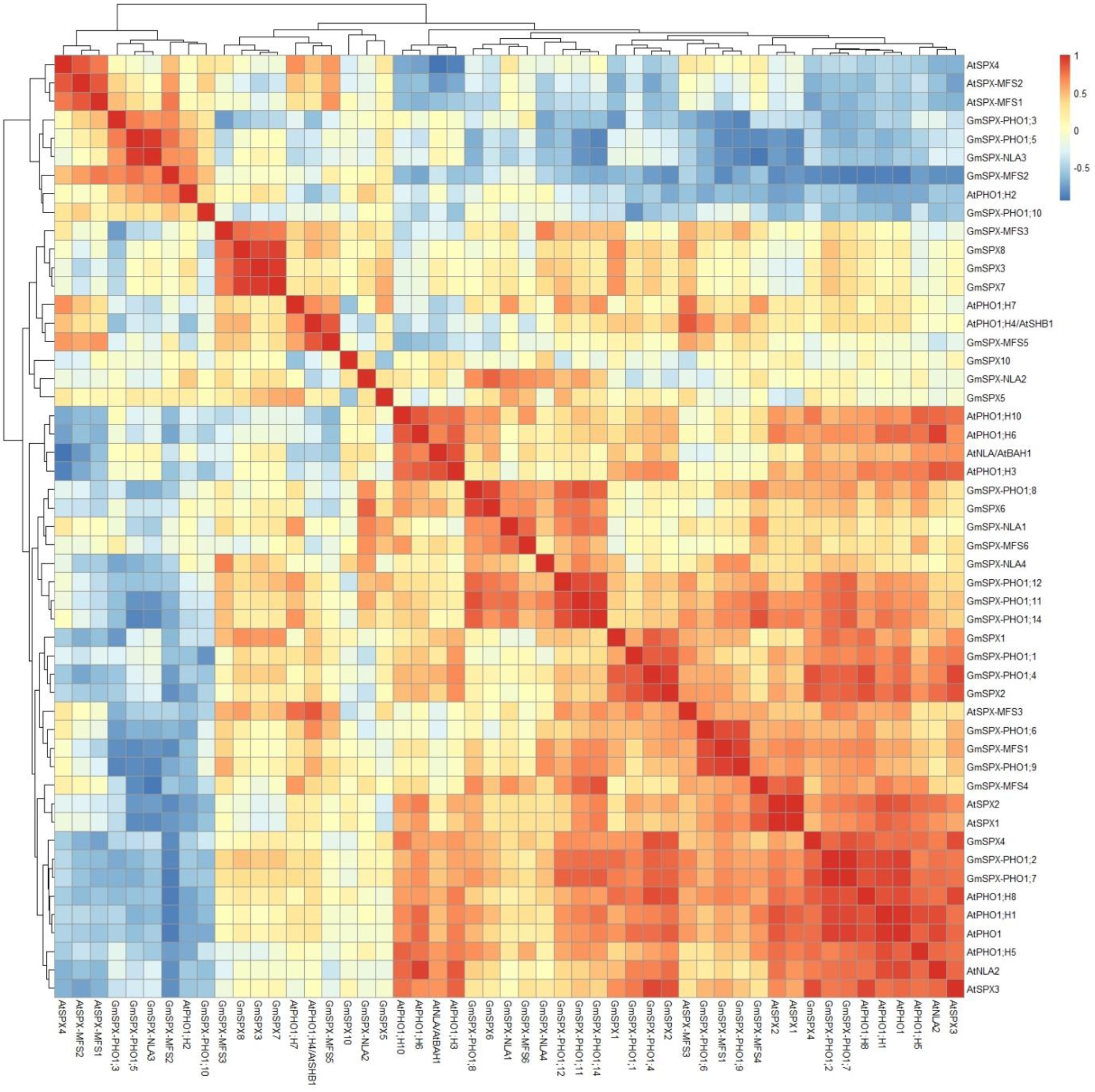
Correlation heat map of SPX genes in soybean and Arabidopsis using RNA-seq datasets from three different zones of roots (GSE64665).

